# Transcriptome-wide dynamics of extensive m6A mRNA methylation during *Plasmodium falciparum* blood-stage development

**DOI:** 10.1101/572891

**Authors:** Sebastian Baumgarten, Jessica M. Bryant, Ameya Sinha, Thibaud Reyser, Peter R. Preiser, Peter C. Dedon, Artur Scherf

## Abstract

Malaria pathogenesis results from the asexual replication of *Plasmodium falciparum* within human red blood cells, which relies on a precisely timed cascade of gene expression over a 48-hour life cycle. Although substantial post-transcriptional regulation of this hardwired program has been observed, it remains unclear how these processes are mediated on a transcriptome-wide level. To this end, we identified mRNA modifications in the *P. falciparum* transcriptome and performed a comprehensive characterization of N^6^-methyladenosine (m^6^A) over the course of blood stage development. Using mass spectrometry and m^6^A RNA sequencing, we demonstrate that m^6^A is highly developmentally regulated, exceeding m^6^A levels known in any other eukaryote. We identify an evolutionarily conserved m^6^A writer complex and show that knockdown of the putative m^6^A methyltransferase by CRISPR interference leads to increased levels of transcripts that normally contain m^6^A. In accordance, we find an inverse correlation between m^6^A status and mRNA stability or translational efficiency. Our data reveal the crucial role of extensive m^6^A mRNA methylation in dynamically fine-tuning the transcriptional program of a unicellular eukaryote as well as a new ‘epitranscriptomic’ layer of gene regulation in malaria parasites.

## Introduction

Malaria, a mosquito-borne human disease caused by the unicellular apicomplexan parasite *Plasmodium falciparum,* remains a major global health threat (World Health Organization, 2018). Human pathogenesis results from the 48-hour intra-erythrocytic developmental cycle (IDC), during which each parasite undergoes schizogony within red blood cells (RBC) to create up to 32 new daughter cells (Fig. 1a). Each step of the IDC – RBC invasion (0-hour post infection [hpi]), host cell remodeling, genome replication, and eventually RBC egress – is effected by a precisely timed cascade of gene expression. Throughout this process, mRNAs from the majority of genes expressed reach peak abundance only once, which is thought to correspond to the time point when its encoded product is most required^1^.

**Fig. 1:**
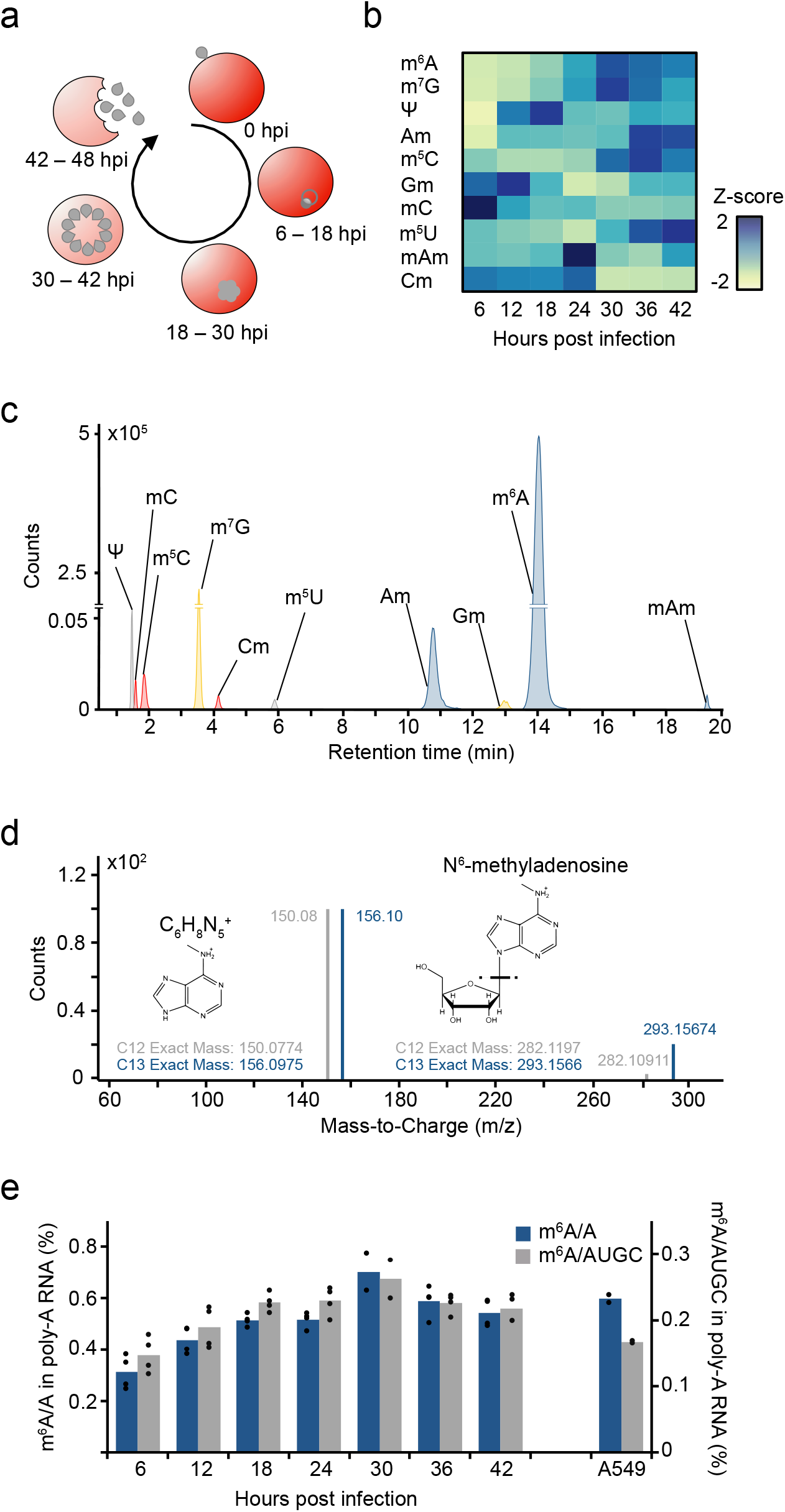
Global dynamics of mRNA modifications in the *P. falciparum* IDC. a: Diagram illustrating the asexual replicative cycle of *P. falciparum* inside human red blood cells including RBC invasion, replication via schizogony, and RBC egress. hpi: hours post infection. b: Heatmap of normalized abundances (modification/canonical base) of ten mRNA modifications measured by LC-MS/MS at six-hour intervals over the course of the IDC. Blue indicates a high Z-score while yellow indicates a low Z-score (see STAR Methods). c: *P. falciparum* mRNA modifications. LC-MS/MS extracted ion chromatograms of modified ribonucleosides analyzed in parasite mRNA collected at 36 hpi. rC: Cytosine, rU: Uracil, rG: Guanosine, rA: Adenosine, m^6^A: N^6^-methyladenosine, m^7^G: N^7^-methyl-2’-guanosine, Ψ: Pseudouridine, Am: 2’-O-methyladenosine, m^5^C: 5-methylcytosine, Gm: 2’-O-methylguanosine, mC: (3 or 4)-methylcytosine, m^5^U: 5-methyluridine, mAm: *-methyl-2’-O-methyladenosine. Cm: 2’-O-methylcytosine. d: Analysis of m^6^A by high-resolution mass spectrometry. Product ion spectrum of m^6^A in *P. falciparum* mRNA (grey) using heavy (C13) labelled *E. coli* tRNA (blue) as a standard. Fragmentation of m/z 282.12 (C12-red) and m/z 293.16 (C13-blue) leads to the breakage of the glycosidic bond resulting in the loss of the ribose sugar (132 Da) to form a daughter ion having a m/z 150.08 (C12-red) and m/z 156.10 (C13-blue). e: Global dynamics of calibrated m^6^A/A (blue) and m^6^A/AUGC (grey) levels across the asexual replicative cycle in six-hour intervals. As a reference, human A549 mRNA was analyzed in parallel^51^. *n* = 4 for all time points, except 30 hpi (n = 2); *n* = 2 for human A549 samples. Points represent individual biological replicates with bars showing the mean.

At the transcriptional level of gene regulation, silencing is achieved via heterochromatinization of genes encoding variant surface antigens^2^ and the inducer of sexual stage development^3^. Heterochromatin, however, is restricted to subtelomeric and several central chromosomal regions^4^, and the large majority of the *P. falciparum* genome remains in a euchromatic and transcriptionally permissive state throughout the IDC^5^. Recent studies of nucleosome occupancy have shown that dynamic chromatin accessibility in promoter regions strongly correlates with mRNA abundance, and that these sequences can be recognized by transcription factors *in vitro*^6–8^. Intriguingly, the parasite’s periodic transcriptional profile appears to be hardwired and unresponsive to perturbation throughout the IDC^9,10^. Therefore, it has been suggested that while transcription may dictate the initial level of mRNA abundance, modulation at the RNA level could provide a more sensitive and/or responsive layer of regulation^11–13^.

Such fine-tuned post-transcriptional regulation in *P. falciparum* is most evident in the dynamics of mRNA degradation rates. mRNA stability was shown to be highly dynamic during the IDC^13^ and to increase gradually over a single developmental cycle^14^. Moreover, extensive post-transcriptional repression of protein synthesis occurs throughout the IDC^15,16^ and translational efficiency was found to differ substantially among mRNA transcripts at the same time point during the IDC^17^. However, the underlying mechanisms orchestrating these processes on a transcriptome-wide scale are as yet unknown.

Several recent studies in model organisms have shown that chemical nucleotide modifications located at internal positions of mRNA can act as mediators of global post-transcriptional regulation of gene expression^18,19^. Most notably, the relatively abundant N^6^-methyladenosine (m^6^A) has been shown to affect a multitude of cellular processes^20^.

In mammalian cells, m^6^A is catalyzed by a methyltransferase complex consisting of a catalytically active m^6^A methyltransferase (methyltransferase-like 13, METTL3), a second methyltransferase-like protein (METTL14) and the METTL3 adaptor protein Wilms-Tumor-1 associated protein (WTAP)^21^. METTL14 does not have enzymatic activity by itself^22^, but through binding of substrate mRNA enhances the activity of METTL3^23^. A key function of WTAP is the localization of the m^6^A writer complex to nuclear speckles^24^. While other proteins can assemble with the m^6^A methyltransferase complex in mammalian cells, orthologs of METTL3, METTL14 and WTAP form a conserved m^6^A core complex that is present in all eukaryotes investigated^21^.

m^6^A influences the fate of individual transcripts by changing the secondary structure of RNAs, making adjacent binding sites accessible for RBPs^25^, or directly recruiting specific m^6^A-binding proteins^21^. In mammalian cells, m^6^A-binding proteins include eukaryotic initiation factor 3 (eIF3) and proteins of the YTH (YT521-B homology) family. m^6^A-mediated recruitment of eIF3 initiates translation independent of the canonical cap-binding eIF4E^26^ whereas YTH proteins primarily decrease mRNA stability in mammalian cells^27,28^. In addition, m^6^A methylation is also involved in alternative splicing and modulating translational efficiencies^29,30^. The cellular and organismal processes affected by m^6^A are widespread and include cell differentiation, cancer cell progression^27,31^ and regulation of the circadian rhythm in mammalian cells^32^, sex determination in *Drosophila*^33^, and meiosis in yeast^34^.

The *P. falciparum* genome has the highest AT content (~80%) of any organism sequenced to date^35^. More intriguing yet is the even stronger adenosine bias in the mRNA transcriptome, with adenosines constituting ~45% of all mRNA bases, compared to ~32% in yeast or ~27% in humans. The reason for this high AT bias remains puzzling, but given this unique nucleotide composition and high level of post-transcriptional regulation during the IDC, we hypothesized that mRNA nucleotide modifications, especially on adenosines, could provide a novel layer of gene regulation in *P. falciparum.*

Here, we use mass spectrometry to identify *P. falciparum* mRNA modifications and show that m^6^A mRNA methylation is an essential layer of post-transcriptional regulation during the IDC. We identify an evolutionarily conserved m^6^A methyltransferase complex, characterize PfMT-A70 as an integral player in m^6^A methylation and show that m^6^A dynamically fine-tunes a hardwired transcriptional program across *P. falciparum* blood-stage development. In addition, this study provides an invaluable resource and the genomic toolkit to explore this new epigenetic regulator in various phases of the complex life cycle of *P. falciparum.*

## Results

### *P. falciparum* intra-erythrocytic development is characterized by an extensive and dynamic m^6^A mRNA methylation program

We first identified mRNA modifications in an unbiased manner and on a transcriptome-wide scale by chromatography-coupled triple quadrupole mass spectrometry (LC-MS/MS) from tightly synchronized parasites harvested at seven time points over the course of the IDC (Supplementary Fig. 1a-c). Ten mRNA modifications that were previously identified in eukaryotic mRNAs were detected in our analysis at varying relative quantities at all time points of the IDC (Fig. 1b,c). These modifications include the putative mRNA cap terminal nucleotide N^7^-methylguanosine (m^7^G), pseudouridine (Ψ), 5-methylcytidine (m^5^C), and N^6^-methyladenosine (m^6^A) (Supplementary Table 1, Fig. 1d, Supplementary Fig. 1d). m^6^A was previously reported to also be the most abundant internal mRNA modification in mammalian cells^36^. While global m^6^A/A levels in *P. falciparum* are comparable to those measured in human mRNA (Fig. 1e, Supplementary Table 2), in combination with the high adenosine content of *P. falciparum* protein-coding mRNA (~45%), the overall m^6^A levels in the parasite’s mRNA exceed those measured in human mRNA almost throughout the IDC (Fig. 1e). Even more strikingly, m^6^A mRNA methylation in *P. falciparum* is highly developmentally regulated, with a continuous increase from 0.3% at early stages of the IDC up to 0.7% m^6^A/A at 30 hpi (Fig. 1e).

A search of the *P. falciparum* genome revealed a highly conserved putative ortholog (PF3D7_0729500) of the catalytically active m^6^A mRNA methyltransferase^22^ characterized in other eukaryotes (i.e. METTL3 in mammalian cells, Ime4 in *Saccharomyces cerevisiae,* and MTA in *Arabidopsis thaliana)* (Fig. 2a). The *P. falciparum* protein contains the characteristic functional domain of its putative orthologs (i.e. MT-A70) and was thus named PfMT-A70. To investigate a potentially conserved role in *P. falciparum,* we generated a cell line expressing an episomal HA-tagged PfMT-A70 protein. In line with the finding in other eukaryotes that m^6^A methylation occurs prior to nuclear mRNA export^37^ we also find that PfMT-A70 localizes to the nucleus and the cytoplasm (Fig. 2b).

**Fig. 2:**
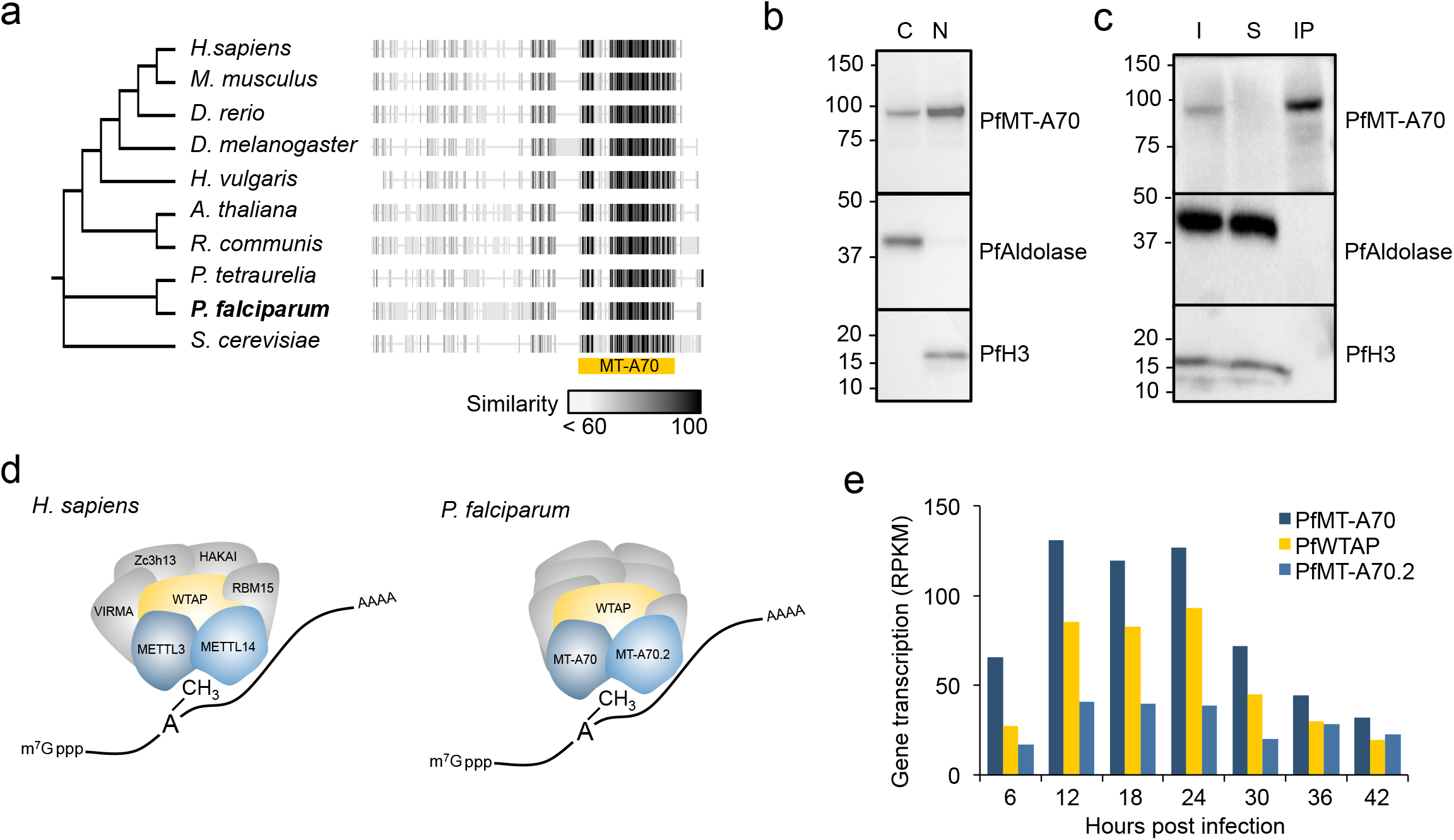
Characterization of the *P. falciparum* m^6^A writer complex. a: Phylogenetic tree and protein alignments of PfMT-A70 with orthologs in other eukaryotes. The location of the MT-A70 domain is shown below the alignment (yellow bar). The grey scale gradient indicates the percentage similarity across all proteins at an aligned position. b: Western blot analysis showing the enrichment of HA-tagged PfMT-A70 in the cytoplasmic (C) and nuclear (N) cell fractions at 12 hpi. PfAldolase and histone H3 serve as controls for the cytoplasmic and nuclear fraction, respectively. Numbers on left indicate molecular weight in kilodaltons. c: Western blot analysis of PfMT-A70-HA co-IP with α-HA antibodies. PfAldolase and histone H3 demonstrate efficacy of the α-HA co-IP. Numbers on left indicate molecular weight in kilodaltons. I: input; S: supernatant; IP: co-IP fraction. d: Schematic of the human m^6^A writer complex (left) and the *P. falciparum* proteins that were co-immunoprecipitated with PfMT-A70. Orthologous proteins are highlighted in the same color (i.e. METTL3/PfMT-A70, METTL14/PfMT-A70.2, WTAP/PfWTAP). Co-factors of the human core m^6^A writer complex (left)^49–52^ and putative, non-conserved co-factors of *P. falciparum* (right) are shown in grey. e: Gene transcription (in reads per kilobase of exon per one million mapped reads [RPKM])^57^ of putative orthologs of the *P. falciparum* m^6^A writer complex at six-hour intervals over the course of the IDC.

Since m^6^A in other eukaryotes is known to be deposited by a multi-protein m^6^A writer complex assembled around the m^6^A mRNA methyltransferase^21^, we next performed protein co-immunoprecipitations (co-IP) of PfMT-A70-HA followed by LC-MS to identify interacting proteins (Fig. 2c,d). In total, we identified 16 proteins that were specifically immunoprecipitated with PfMT-A70 (Supplementary Table 3). Most importantly, among those we find one (PF3D7_1230800) that is highly conserved with eukaryotic orthologs of WTAP (Fig. 2d, Supplementary Fig. 2a). The *P. falciparum* ortholog similarly contains a WTAP/MUM2 domain and was thus named PfWTAP. We also identified a second putative mRNA methyltransferase containing an MT-A70 domain (PF3D7_1235500, named PfMT-A70.2) that shows high sequence conservation with orthologs of the bilaterian METTL14 (Fig. 2d, Supplementary Fig. 2a). All three *P. falciparum* orthologs reach peak transcription levels between 12 and 24 hpi, shortly before maximal m^6^A/A ratios are reached (Fig. 1e, 2e). Among the other co-immunoprecipitated proteins are known RNA/DNA binding proteins such as Alba3, an ATP-dependent RNA helicase, and a FoP-domain containing protein as well as several of unknown function (Supplementary Table 3).

### CRISPR interference of PfMT-A70 decreases global m^6^A mRNA methylation

We next targeted PfMT-A70 to gain further insight into the functional role of m^6^A mRNA methylation in *P. falciparum.* However, attempted CRISPR/Cas9-mediated deletion of the C-terminal catalytic MT-A70 domain was unsuccessful, suggesting that PfMT-A70 is essential for survival. In support of our finding, a recent random mutagenesis study failed to mutagenize this locus^38^. Similarly, C-terminal tagging of the protein with a ligand-regulated *Escherichia coli* dihydrofolate reductase destabilization domain^39^ for protein knockdown was unsuccessful. We therefore applied a genome editing-free method for gene down-regulation termed CRISPR interference (CRISPRi)^40^ which has recently been adapted for *Plasmodium*^41,42^ and is based on an enzymatically inactive (‘dead’) Cas9 protein (dCas9). In this method, the Cas9 protein is rendered inactive by two point mutations (Supplementary Fig. 2b)^40,43^, allowing it to bind to, but not cut, its target locus via a specific single-guide RNA (gRNA). This binding interferes with the transcription machinery and leads to down-regulation of gene transcription. For CRISPRi of PfMT-A70, we targeted the PfMT-A70 promoter downstream of the transcription start site and ~100 bp upstream of the translation start site with a gRNA complementary to the non-template strand (‘gPfMT-A70’, Fig. 3a). A second cell line expressing a non-specific gRNA was used as a negative control (‘gControl’, Fig. 3a).

**Fig. 3:**
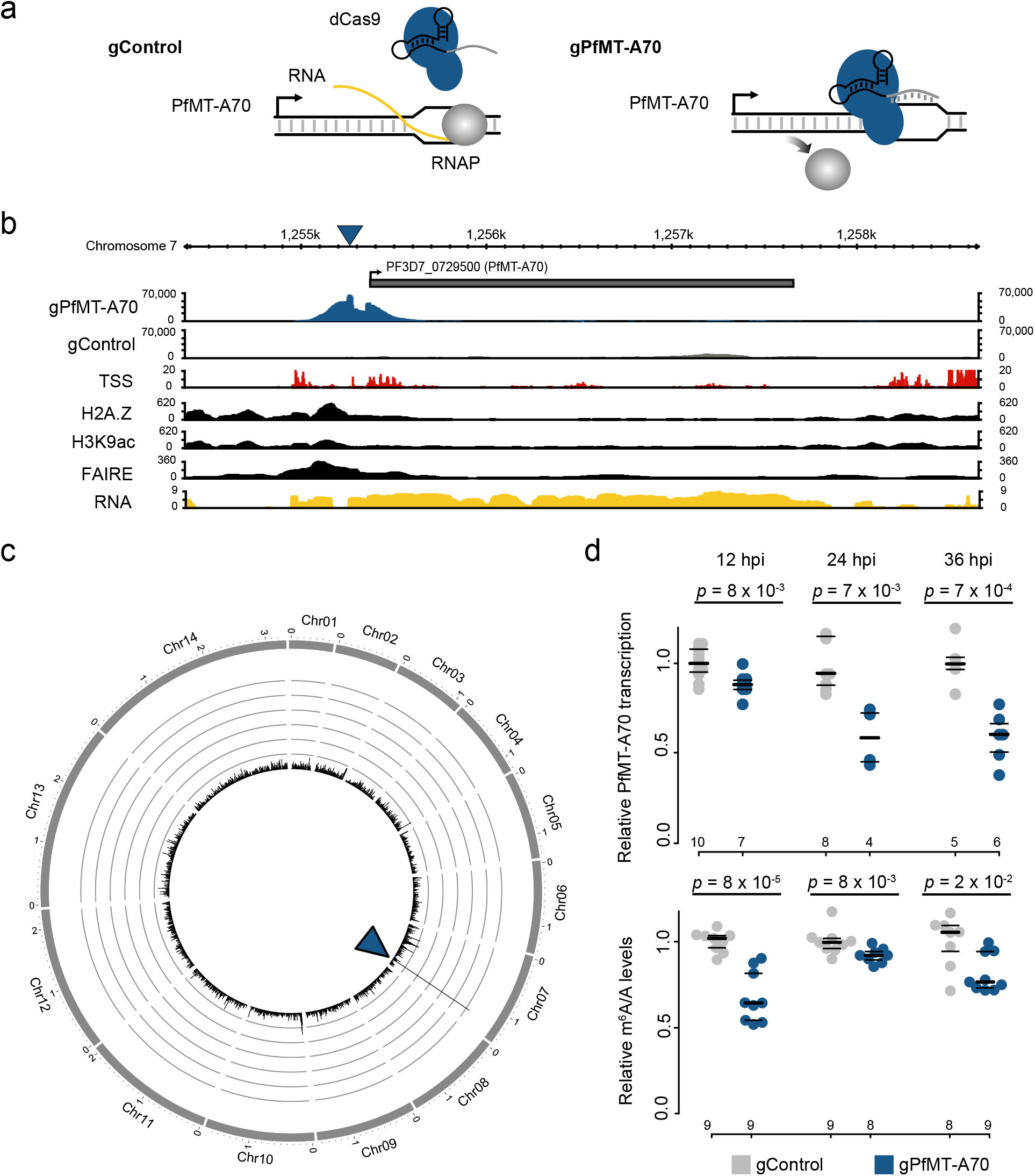
Knockdown of the PfMT-A70 m^6^A methyltransferase by CRISPR interference. a: Diagram of the CRISPR interference system for targeted transcriptional knockdown of PfMT-A70. A non-specific gRNA (gControl) without binding site in the *P. falciparum* genome is used as a negative control in all experiments (left). The specific gRNA (gPfMT-A70) targets the PfMT-A70 promoter on the non-template strand downstream of the putative transcription start site, blocking the elongating RNA polymerase II (RNAP) and silencing the target gene (right). b: dCas9 ChIP sequencing data (12 hpi) show enrichment for gPfMT-A70-targeted dCas9 (blue), but not non-targeted dCas9 (gControl in grey) at the targeted upstream region of the PfMT-A70 gene (indicated at top with a grey bar). Arrow indicates direction of transcription. Blue triangle indicates location of the gRNA target site. Genomic features investigated in previous studies that define the putative promoter region are shown below: transcription start site (TSS)^46^, histone 2A variant (H2A.Z)^45^, acetylation of histone H3 at lysine 9 (H3K9ac)^5^, nucleosome depletion as determined by FAIRE-seq^7^ (FAIRE), and RNA-seq coverage (RNA) at the genomic locus^6^. c: Circos plot showing dCas9 ChIP-seq data for gPfMT-A70-targeted dCas9 across all 14 nuclear chromosomes (exterior grey bars), normalized to gControl. The arrowhead indicates the specific target site for gPfMT-A70. Chromosome name and length (in megabases) indicated at the exterior of the plot. Y-axis indicates ChIP enrichment on a scale from 0 to 150 RPKM in 1000 nt windows. d: Comparison of PfMT-A70 transcription measured by RT-qPCR (top) and m^6^A/A measured by LC-MS/MS (bottom) between gControl (grey) and gPfMT-A70 (blue) parasites at three different time points over the IDC (indicated at the top). PfMT-A70 transcript levels were normalized to those of the housekeeping gene serine tRNA ligase (PF3D7_0717700). The number of biological replicates is indicated on the bottom of the graph. p-values were calculated using two-tailed independent-samples t-test (RT-qPCR) and a Mann-Whitney U test (LC-MS/MS). Average of PfMT-A70 transcription and m^6^A/A levels in gControl parasites were set to 1. Horizontal lines represent median and interquartile range.

To validate the specific binding of dCas9 in gPfMT-A70 parasites, we adapted a protocol for dCas9-associated chromatin immunoprecipitation followed by sequencing (dCas9 ChIP-seq)^44^. We found a robust enrichment of gPfMT-A70-directed dCas9 at the target site, which was not seen for the control gRNA (Fig. 3b). Furthermore, the gPfMT-A70 dCas9 footprint overlaps with a nucleosome-depleted region (FAIRE)^7^ enriched for H3K9ac^5^ and H2A.Z^45^, which are all hallmarks of promoter regions in *P. falciparum*^46^ (Fig. 3b). A genome-wide analysis for off-target binding events of gPfMT-A70 dCas9 showed no substantial enrichment other than the PfMT-A70 promoter, demonstrating the high specificity of the method (Fig. 3c).

To assess the effect of CRISPRi on PfMT-A70 expression and m^6^A/A ratios across the *P. falciparum* IDC, we harvested mRNA from tightly synchronized parasites for both the gPfMT-A70 and gControl cell line at 12, 24, and 36 hpi. Transcription levels of PfMT-A70 measured by RT-qPCR (reverse transcription quantitative polymerase chain reaction) revealed a significant down-regulation of the target gene throughout the IDC, with levels of knockdown ranging from ~ 15% at 12 hpi to 40% at 24 and 36 hpi (Fig. 3d, top). Although CRISPRi of PfMT-A70 did not show a discernible effect on parasite growth (Supplementary Fig. 2c), subsequent measurements of global m^6^A/A levels by LC-MS/MS revealed a significant 10-30% decrease at all three time points, (Fig. 3d, bottom), further identifying PfMT-A70 as an integral component of the m^6^A writer complex in *P. falciparum.*

### mRNA transcripts are differentially m^6^A-methylated throughout the IDC

We next assessed the dynamics of m^6^A mRNA methylation on a transcript-specific level across the IDC of *P. falciparum* using m^6^A-methylated mRNA immunoprecipitation (IP) followed by sequencing (m^6^A-seq) (Supplementary Fig. 3a,b). An initial assessment of m^6^A antibody specificity for *P. falciparum* m^6^A-methylated mRNA by LC-MS/MS showed a consistent m^6^A enrichment and depletion in the m^6^A-IP fraction and the supernatant, respectively (Fig. 4a, Supplementary Fig. 3c). We next prepared directional RNA sequencing libraries from m^6^A eluates (‘m^6^A-IP’) and mRNA input (‘m^6^A-input’, used for peak normalization) from two replicates of gPfMT-A70 and gControl parasites at 12, 24, and 36 hpi. Stringent read mapping and filtering showed that > 99% of all reads from the m^6^A-IP map to protein-coding regions, confirming that the m^6^A LC-MS/MS measurements result from mRNA transcript and not ribosomal or transfer RNAs.

**Fig. 4:**
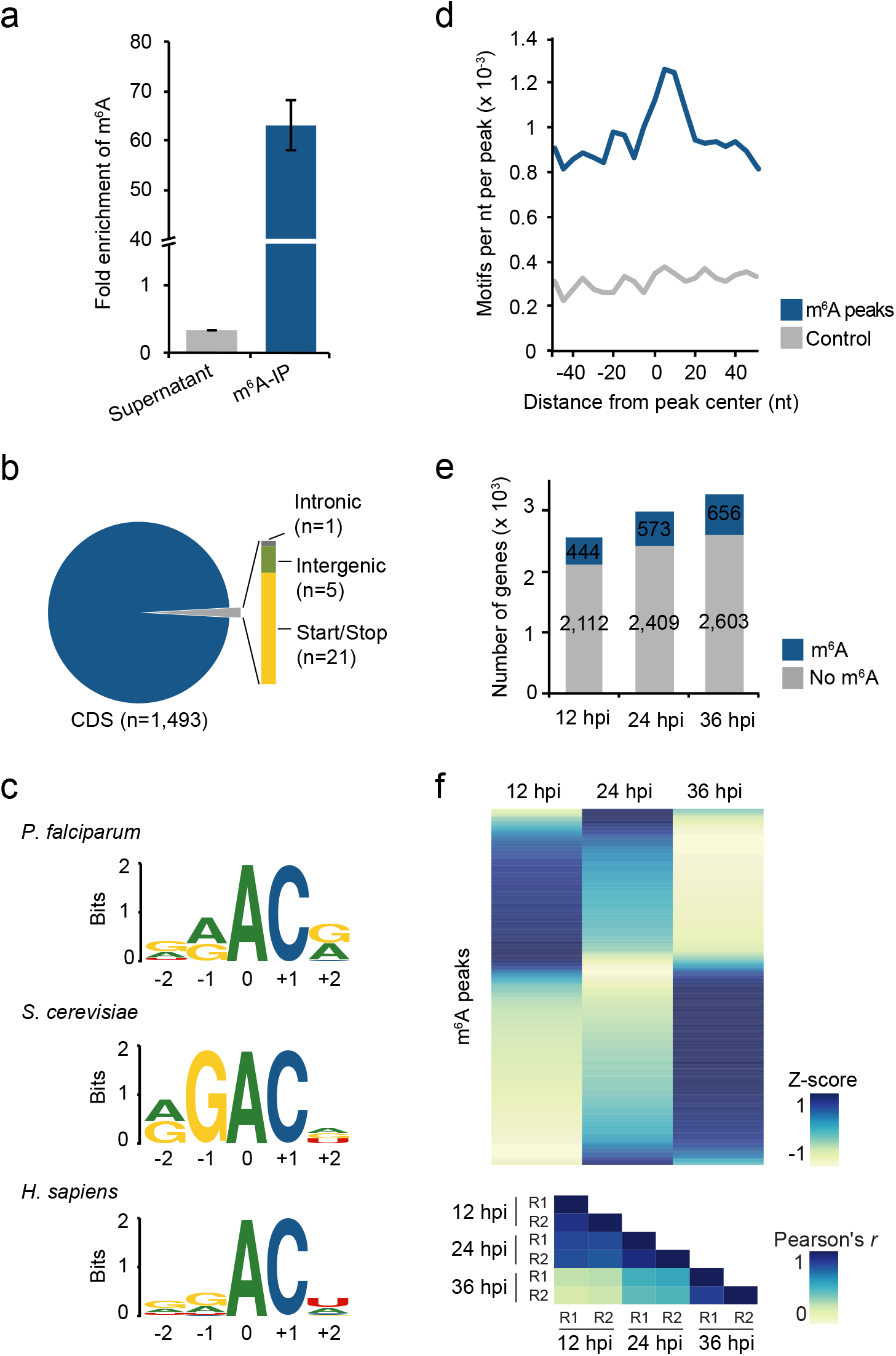
Differential m^6^A methylation during the *P. falciparum* IDC. a: Fold enrichment of m^6^A levels as measured by LC-MS/MS in different fractions of the m^6^A-IP experiment, showing a three-fold depletion in the supernatant and 63-fold enrichment in the eluate. m^6^A levels were normalized to the total number of adenosines (i.e. m^6^A/A) for each fraction and fold enrichments were calculated as ratios over the m^6^A/A levels in the input mRNA sample (collected at 36 hpi). Error bars indicate standard error of the mean for two replicates. b: Distribution of all significant m^6^A-seq peaks (FDR ≤ 0.05) in different genomic regions: within introns (Intronic), outside of coding regions (Intergenic), inside coding sequences (CDS), within 200 nt upstream of the translation start or downstream of the translation stop site (Start/Stop). The total number of identified m^6^A sites by m^6^A-seq is likely an underestimate. Besides those that were not called due to our conservative annotation approach, m^6^A sites in transcripts reaching peak expression at different time points or peaks in low complexity regions have likely been missed c: Comparison of the deduced sequence motif of m^6^A methylation sites in *P. falciparum* (top), S. *cerevisiae* (middle)^34^, and human (bottom)^47^. The relative height of each nucleotide indicates frequency and the total height at each position represents sequence conservation (i.e. ‘bits’). The N^6^-methylated adenosine is located at position 0. d: Density plot showing the occurrence of the consensus motif within ± 50 nt of all significant m^6^A-seq peak summits (blue) and size-matched random control sequences (grey) in 5 nt windows. e: Bar charts indicating the number of transcripts (RPKM ≥ 5) with no (grey) or at least one significant (blue) m^6^A site at three different time points over the IDC. f: Heatmap of average row Z-score normalized m^6^A enrichments in gControl parasites displaying changes in transcript-specific methylation at three time points throughout the IDC (top). Each row represents an m^6^A peak in a transcript expressed with RPKM ≥ 5 at all three time points (n = 840). Bottom: Heatmap showing Pearson’s *r* correlation coefficients of m^6^A enrichments, demonstrating a high degree of reproducibility between the two replicates (R1, R2) at each time point. Blue indicates a high score while yellow indicates a low score.

To map m^6^A methylation sites, we called m^6^A peaks separately for each replicate of gPfMT-A70 and gControl samples collected at every time point. Sites were only considered if they were identified as significantly enriched (FDR ≤ 0.05) in at least two out of four samples collected at a given time point. This approach identified 603, 873, and 996 significant m^6^A peaks at 12, 24, and 36 hpi, a pattern that parallels the global increase in m^6^A/A levels as measured by LC-MS/MS. At all three time points, the majority of transcripts contain only one m^6^A peak (Supplementary Fig. 3d), with an average of ~0.7 m^6^A peaks per kilobase of exon (Supplementary Fig. 3e). In total, we identified 1,520 non-overlapping m^6^A peaks across all time points, present in 1,043 distinct gene transcripts (Supplementary Table 4-6). The large majority of identified m^6^A peak summits in *P. falciparum* are found within protein-coding sequences and in a region 200 nucleotides (nt) upstream of the translation start site or downstream of the translation stop site (Fig. 4b). Only five m^6^A peaks were identified in intergenic regions and only one significant m^6^A peak was found in an intron.

We searched for sequence motifs associated with methylation sites in the ± 100 nt surrounding the summit of the m^6^A peak and identified a single significantly enriched GGACA motif (p = 1 x 10^-12^). This motif is identical to the most enriched 5-mer identified in m^6^A sites of S. *cerevisiae*^34^ and similar to the DRACH (D = G/A/U; R = G/A; H = C/A/U) consensus motif of mammalian m^6^A sites^21^. In addition, we find that motifs in an RAC context are generally enriched compared to a control sequence set of similar length and size (Supplementary Fig. 3f), with the deduced consensus motif being similar to those of S. *cerevisiae* and human m^6^A methylation sites (Fig. 4c)^34,47^. This motif is not only more abundant in m^6^A peaks but is also centrally enriched in a region ± 20 nt around the m^6^A peak summit (Fig. 4d), suggesting a certain context dependency for specific m^6^A methylation sites.

For an inherently dynamic transcriptional program such as that underlying the IDC of *P. falciparum,* changes in transcript-specific methylation might provide a means of modulating the transcriptome independently of initial gene transcription. Throughout the IDC, there is a modest increase in the total number of transcribed genes in gControl parasites that have at least one m^6^A peak (Fig. 4e). However, the m^6^A enrichment per gene, or fraction of transcripts from one gene containing a specific m^6^A site (i.e. m^6^A-IP/m^6^A-input, ‘m^6^A enrichment’), changes extensively throughout the IDC (Fig. 4f). Most m^6^A peaks reach maximum m^6^A enrichment at only one time point, suggesting that transcript-specific methylation is indeed an actively regulated mechanism.

### m^6^A inversely correlates with mRNA stability and translational efficiency

To determine the effect of m^6^A on individual transcripts, we first compared m^6^A enrichment of individual m^6^A peaks between gControl and gPfMT-A70 parasites and found a significant reduction in the gPfMT-A70 cell line (Fig. 5a,b top) at all three time points, mirroring the decrease in global m^6^A/A levels in gPfMT-A70 parasites as measured by LC-MS/MS (Fig. 3d, bottom). These decreases are consistent, with two replicates each of gControl and gPfMT-A70 showing a high degree of correlation at each time point (Fig. 4b, bottom, Supplementary Table 4-S6).

**Fig. 5:**
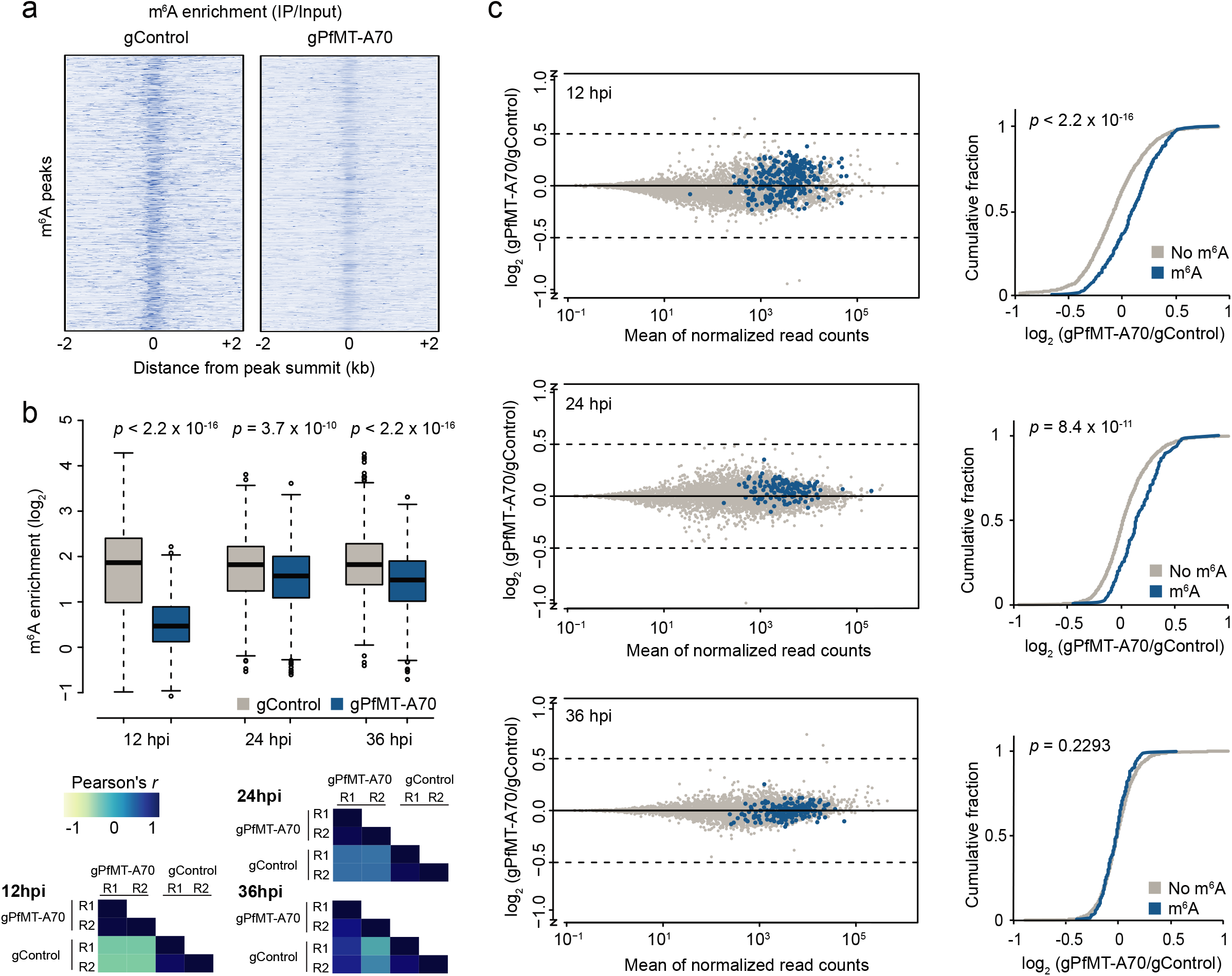
PfMT-A70 knockdown leads to upregulation of m^6^A-methylated transcripts. a: Heatmap of row Z-score normalized m^6^A enrichment at 12 hpi depicting the decrease in m^6^A enrichment for individual m^6^A peaks (n = 603, y-axis) between gControl (left) and gPfMT-A70 (right) parasites calculated in 10 nt windows in a region ± 2 kb around m^6^A peak summits. Darker color indicates higher enrichment. b: Boxplot of m^6^A enrichment for all m^6^A peaks in gControl (grey) and gPfMT-A70 (blue) parasites showing a significant decrease in m^6^A enrichment following PfMT-A70 depletion at all three time points. m^6^A enrichment for each peak was averaged over two replicates in each condition and time point (see correlation between replicates below). p-values were calculated using a Mann-Whitney U test. Bottom: Heatmap depicting Pearson’s *r* correlation coefficients of m^6^A enrichment among replicates of gControl and gPfMT-A70 parasites at all three time points (see Supplementary Table 4-S6). Blue indicates a high score while yellow indicates a low score. c: MA-plots (left) of log_2_ fold changes (gPfMT-A70/gControl, M) plotted over the mean abundance of each gene *(A)* at 12 (top), 24 (middle) and 36 hpi (bottom). n = 3 for gPfMT-A70 and n = 2 for gControl at all time points. Transcripts containing m^6^A peaks with a more than two-fold decrease in m^6^A enrichment following PfMT-A70 depletion are highlighted in blue (12 hpi: n = 284; 24 hpi: n = 132; 36 hpi n = 199). Cumulative fraction plots (right) comparing the distribution of log_2_ fold-changes (gPfMT-A70/gControl) between non-methylated transcripts (grey) and transcripts with a more than two-fold decrease in m^6^A enrichment following PfMT-A70 depletion (blue) at 12 (top), 24 (middle) and 36 hpi (bottom). *p-values* were calculated using a Mann-Whitney U test.

We next performed directional mRNA sequencing and compared transcript abundances between gControl and gPfMT-A70 parasites at 12, 24, and 36 hpi. At all time points, we observed minimal changes in global transcript abundance, with only a small percentage of genes showing significant differential expression (Supplementary Fig. 4a, Supplementary Table 7). In contrast, at 12 and 24 hpi, we found a significant increase in overall abundance of transcripts that show the most pronounced decrease in m^6^A enrichment (≥ two-fold) upon PfMT-A70 depletion (Fig. 5c).

Theoretically, this observed increase in transcript abundance following PfMT-A70 knockdown is due either to higher transcription rates or lower transcript degradation rates. Because m^6^A methylation was previously shown to affect mRNA degradation^27,28^, we compared mRNA stability data (measured as mRNA half-life) previously obtained from wild type parasites^14^ between m^6^A-methylated and non-methylated transcripts found in our study. Strikingly, m^6^A-methylated transcripts have significantly lower mRNA stabilities than non-methylated transcripts (Fig. 6a, left). These differences are significant at 12 (p = 5.3 x 10^-5^) and 24 hpi (p = 9.1 x 10^-9^), but not at the late 36 hpi time point (p = 0.59). In addition, we find even lower mRNA stabilities for the ‘m^6^A-sensitive’ transcripts (i.e. ≥ two-fold decrease in m^6^A enrichment and increased transcript abundance [log_2_ fold-change > 0]) at 12 and 24 hpi (Fig. 6a, right). These data suggest that the increase of transcript abundances following PfMT-A70 knockdown might indeed reflect decreased rates of m^6^A-mediated mRNA degradation at both the 12 and 24 hpi time point. A further comparison of m^6^A methylation with additional mRNA stability data^13^ revealed a similar pattern, with m^6^A-methylated transcripts having significantly lower mRNA stabilities than non-methylated transcripts (Supplementary Fig. 4b).

**Fig. 6:**
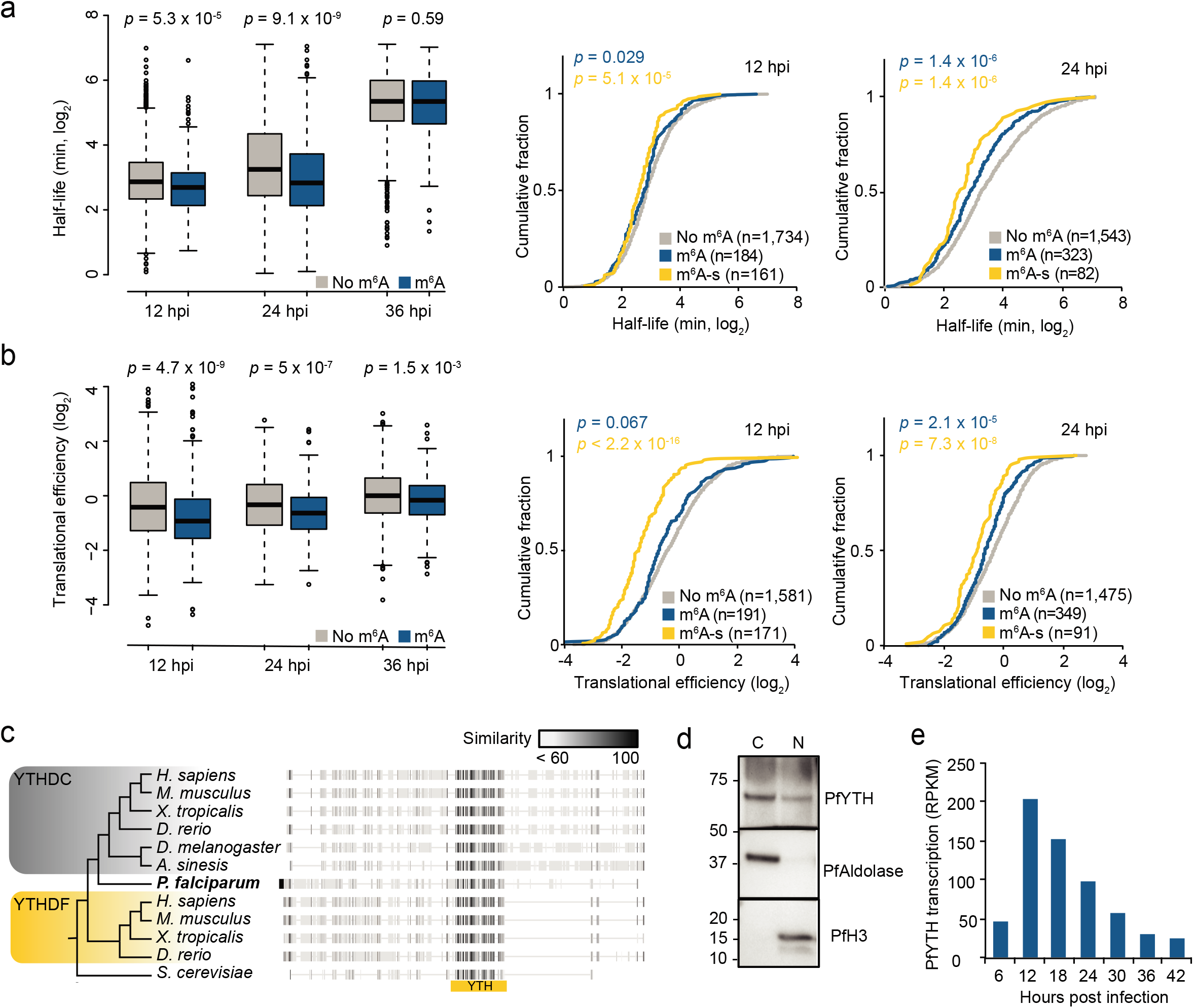
m^6^A methylation inversely correlates with mRNA stability and translational efficiency. a and b: Left: Boxplots depicting mRNA half-lives (a) or translation efficiency (b) for transcripts without (grey) or with (blue) m^6^A methylation sites at 12, 24, and 36 hpi. *p-values* were calculated with a Mann-Whitney U test. Right: Cumulative fraction plots of mRNA half-lives (a) and translation efficiency (b) at 12 and 24 hpi for transcripts without m^6^A site (grey), with m^6^A sites (blue), and ‘m^6^A-sensitive’ transcripts (yellow). p-values were calculated with a Mann-Whitney U test against mRNA half-lives or translation efficiencies of non-methylated transcripts. c: Phylogenetic tree and protein alignment of the putative m^6^A reader protein PfYTH and orthologs in other eukaryotes. The location of the YTH domain is indicated below (yellow bar). The grey scale gradient indicates the percentage similarity across all proteins at an aligned position. YTHDC: YTH domain-containing protein; YTHDF: YTH domain-containing family protein. d: Western blot analysis showing the enrichment of HA-tagged PfYTH in the cytoplasmic (C) and nuclear (N) fractions at 12 hpi. PfAldolase and histone H3 serve as controls for the cytoplasmic and nuclear fractions, respectively. Numbers on left indicate molecular weight in kilodaltons. e: Transcription (RPKM) of the *P. falciparum* PfYTH protein over the course of the IDC^57^.

We next compared translational efficiencies (TE) between m^6^A-methylated and non-methylated transcripts throughout the IDC. TEs were calculated previously for *P. falciparum* parasites as the ratio of ribosome-bound RNA versus steady state mRNA 17. At each developmental stage, m^6^A-methylation correlates with significantly lower translational efficiencies for a transcript (Fig. 6b, left). This correlation is most pronounced at the 12 (*p* = 4.7 x 10^-9^) and 24 hpi (*p* = 5 x 10^-7^) time points, but is still significant at 36 hpi (*p* = 1.5 x 10^-3^). Furthermore, similar to the pattern seen for mRNA stability, we observe an additional significant decrease in translational efficiencies between m^6^A methylated and the subset of ‘m^6^A-sensitive’ transcripts at 12 and 24 hpi (Fig. 6b, right).

The ‘m^6^A-sensitive’ transcripts are enriched in genes involved in biological processes that are highly relevant for their respective time points of development (Supplementary Fig. 4c). At 12 hpi, these processes include regulation of gene expression (*q* = 3 x 10^-3^) and DNA-template transcription and regulation of RNA biosynthesis (*q* = 3 x 10^-3^). Among these gene transcripts are four members of the AP2 transcription factor family, two histone deacetylases, one histone-lysine methyltransferase, and one bromodomain-containing protein (Supplementary Table 8). Similarly, at 24 hpi, binding to chromatin (*q* = 6.1 x 10^-3^) and protein binding are among the most significantly enriched molecular functions. Gene transcripts included in these categories include the high mobility group protein B3, transcriptional coactivator ADA2, SNF2 helicases, two SET domain-containing proteins, one bromodomain-containing protein, and pre-mRNA splicing factors (Supplementary Table 8).

## Discussion

In this study, we characterized dynamic and extensive m^6^A mRNA methylation during the blood stage development of the human malaria parasite. We show that this mRNA modification is developmentally regulated and fine-tunes an otherwise hardwired transcriptional cascade during the asexual life cycle of *P. falciparum.* We generated a high-resolution map of m^6^A methylation sites throughout the IDC and identify PfMT-A70 as an essential member of the putative m^6^A-methyltransferase complex.

As gene knockout was not possible, we used the recently developed CRISPRi system^41–43^ to demonstrate the role of PfMT-A70 in the m^6^A methylation pathway. Because *P. falciparum* lacks the RNA interference pathway, application of CRISPRi is an invaluable new genome editing-free technique for targeted gene knockdown and functional characterization of essential genes in the parasite. Indeed, the CRISPRi strategy allowed us to reduce PfMT-A70 transcript levels to the extent that a consistent phenotype of decreased m^6^A levels via LC-MS/MS and m^6^A-seq, as well as increased abundance of transcripts with decreased m^6^A-enrichment could be observed. Core features of the *P. falciparum* m^6^A methylation machinery are evolutionarily conserved, especially the m^6^A-methyltransferase, PfMT-A70, and the core members of the m^6^A writer complex, PfMT-A70.2 and PfWTAP. Our purification of this complex demonstrates the deep evolutionary origin of m^6^A methylation and highlights its importance in eukaryotic post-transcriptional regulation. In line with this conservation, m^6^A methylation in *P. falciparum* also occurs primarily in an RAC context (R = A/G), which is similar to the m^6^A nucleotide context identified in yeast (RGAC; R = A/G)^34^ and mammalian cells (DRACH; D = G/A/U, R = A/G; H = A/C/U)^48^.

In mammalian cells, several adaptor proteins can assemble with and guide the core m^6^A writer complex^49–52^. However, we did not identify orthologs of these cofactors among the proteins that specifically co-immunoprecipitated with PfMT-A70. It is possible that the putative *P. falciparum* co-factors are lineage-specific, as several of them do not share any sequence homology to other eukaryotic proteins. Most of the co-factors we identified show nuclear localization, and PfMT-A70.2 has been found exclusively in the nuclear proteome^53^. These data suggest that m^6^A methylation in *P. falciparum* occurs prior to nuclear mRNA export^37^. Future functional studies will define the roles of each of these co-factors and the core proteins in the m^6^A writer complex.

Other than the apparent conservation of the core m^6^A writer complex and m^6^A site context, the *P. falciparum* m^6^A methylation program is set apart from other eukaryotes studied by several key aspects. With the high adenosine content in the parasite’s protein-coding transcriptome, overall m^6^A levels are exceptionally high and markedly exceed those measured in human mRNA, suggesting a key role for this modification during asexual replication. It is tempting to speculate that m^6^A methylation might represent one evolutionary driving force of high adenosine contents observed at gene loci. The essentiality of PfMT-A70 and the functional diversity of transcripts containing m^6^A further suggest that this modification is involved in various important cellular processes during the IDC. While m^6^A may directly affect the fate of an mRNA transcript and its encoded protein, the modulation of transcripts encoding major regulatory proteins such as AP2 transcription factors or chromatin remodelers might further amplify its modulatory potential.

In addition to the high levels of m^6^A methylation in *P. falciparum,* the dynamic nature of this modification is also striking. Successive LC-MS/MS measurements and transcript-specific m^6^A-seq independently demonstrated similar m^6^A dynamics with global m^6^A/A levels and the number of identified m^6^A peaks gradually increasing towards the end of the IDC. Dynamic m^6^A methylation is generally controlled through the specific spatio-temporal expression of the m^6^A writer complex, for example in a defined tissue or cell type^33^ or in response to certain conditions^34^. In contrast, we observe dramatic changes in the percentage of methylated transcripts originating from one gene over the course of the *P. falciparum* IDC. In mammalian systems, it was shown that in general, m^6^A is deposited early and maintained throughout the life of an mRNA transcript^37^, although the m^6^A RNA-demethylase Alkbh5 may modulate methylation levels in some cases^56^. Since we did not find a potential m^6^A demethylase in the *P. falciparum* genome, m^6^A-methylation dynamics during the IDC could result from different rates of active m^6^A-methylation by the writer complex and the subsequent rate of m^6^A-methylated transcript degradation. Indeed, the observed gradual increase of global m^6^A/A levels measured by LC-MS/MS might be partially due to the m^6^A-mediated mRNA degradation at earlier but not later stages of the IDC, as is evidenced by the inverse correlation of m^6^A status and mRNA stability at 12 and 24, but not 36 hpi. This is further corroborated by our finding that transcripts with decreased m^6^A enrichment are overabundant only at the earlier, but not the late time point following PfMT-A70 depletion. This discrepancy in m^6^A-dependent mRNA degradation rates might be due to the presence or absence of a specific m^6^A reader protein. In mammalian systems, proteins of the YT521-B homology (YTH) family have been identified as specific m^6^A ‘readers’ that mediate mRNA transcript stability^28^. We also identified a single putative YTH ortholog in *P. falciparum* (PfYTH, Fig. 6c) that is present in both the nucleus and cytoplasm (Fig. 6d). PfYTH transcription decreases over the course of the IDC (Fig. 6e), thus providing a potential link between the gradual increase in global m^6^A/A levels and the decrease in m^6^A-mediated transcript degradation during parasite development.

The major factor controlling mRNA transcript abundance during the IDC is certainly the transcriptional activity at the genic locus^8,13,54,55^. However, here we show that the fate of individual transcripts from the same gene can be changed, possibly by decreasing mRNA stability and/or repressing translation through specific m^6^A methylation. Our findings that 1) transcripts with decreased m^6^A-enrichment following PfMT-A70 knockdown were consistently more abundant and 2) there is an increased association between ‘m^6^A-sensitivity’ and mRNA stability and translational repression suggest that m^6^A serves as a balance to modulate the outcome of protein synthesis independent of initial transcriptional rates. Thus, temporally dynamic changes of transcript-specific m^6^A methylation rates during the IDC have, to a certain extent, the potential to fine-tune an otherwise imprecise, hardwired transcriptional program. Collectively, our study unravels a new layer of dynamic and widespread post-transcriptional modulation of gene expression in *P. falciparum.* The conservation of core features of m^6^A mRNA methylation makes *P. falciparum* an excellent system to study the interplay between m^6^A methylation and gene transcription. Moreover, our results add m^6^A as a major player in the malaria parasite ‘epitranscriptome’ code and open new avenues for drug development in malaria parasites.

## Acknowledgments

We thank Aurelie Claës, Christine Scheidig-Benatar and Patty Chen for help with parasite culture. Protein mass-spectrometry was performed at the Biopolymers and Proteomics core of The Koch Institute Swanson Biotechnology Center. This work was supported by a European Research Council Advanced Grant (PlasmoSilencing 670301) and the French Parasitology consortium ParaFrap (ANR-11-LABX0024) to A.Scherf. Work in the labs of P.R.P. and P.C.D. was funded by the National Research Foundation Singapore under its Singapore-MIT Alliance for Research and Technology (SMART) Centre, Infectious Disease and Antimicrobial Resistance IRGs. S.B. and J.M.B. were supported by an EMBO fellowship (S.B.: ALTF 1444-2016; J.M.B.: ALTF 180-2015). A.Sinha acknowledges financial support from the Singapore-MIT Alliance (SMA) Graduate Fellowships.

## Author contribution

P.R.P, P.C.D. and A.Scherf conceptualized the project. S.B., J.M.B. and A.Scherf conceived experiments. J.M.B. developed and performed CRISPR interference and dCas9 ChIP-seq experiments. S.B. performed m^6^A-seq and RT-qPCR experiments. A.Sinha performed and analyzed LC-MS/MS and protein co-IP experiments. S.B., J.M.B. and T.R. generated constructs, transfectants and parasite material. S.B. performed bioinformatic analyses. P.R.P., P.C.D. and A.Scherf supervised and helped interpret analyses. All authors discussed and approved the manuscript.

---

**Supplementary Fig. 1:**
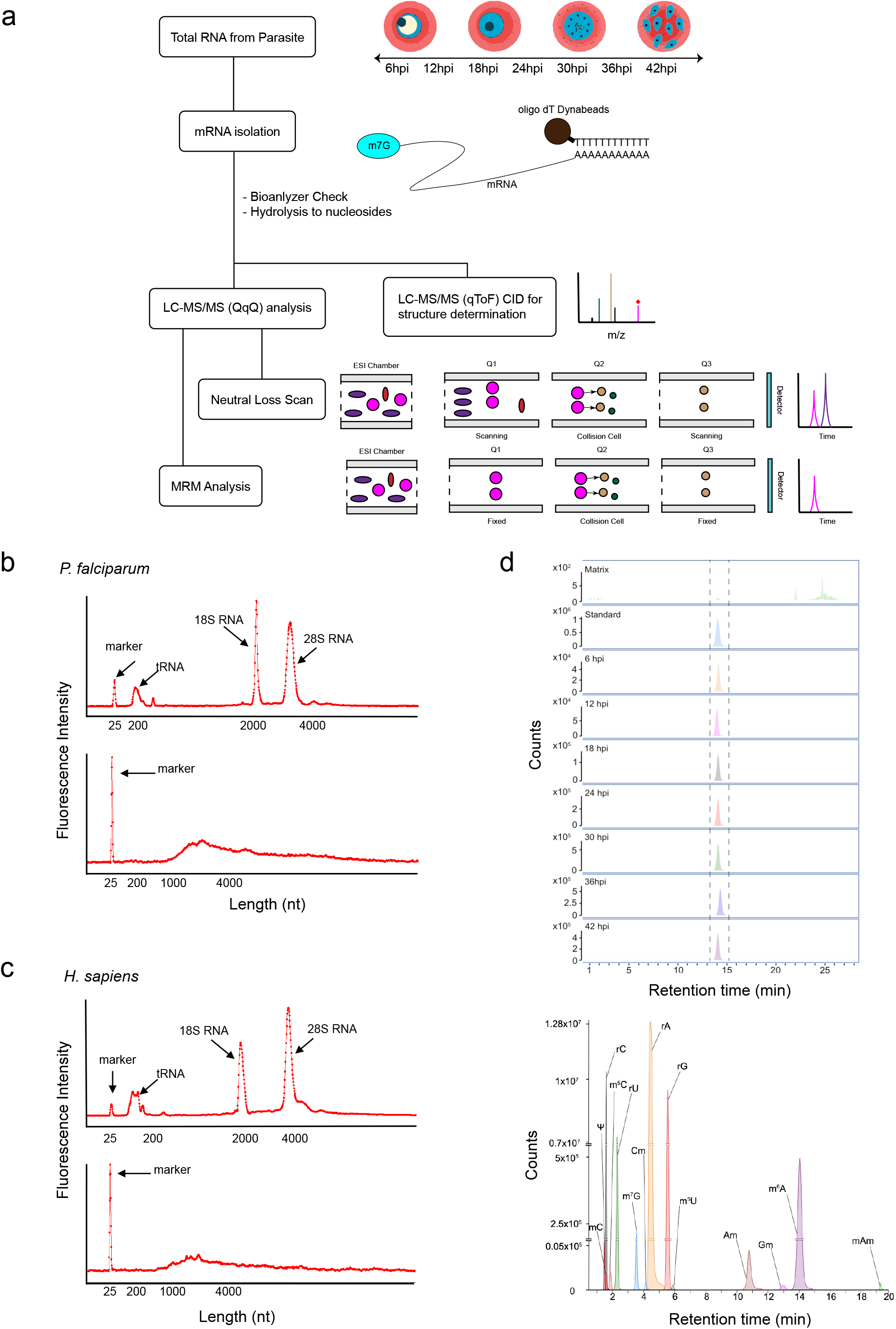
LC-MS/MS measurements of *P. falciparum* mRNA modifications. a: Approach for identification of mRNA modifications in *P. falciparum.* The mRNA samples were isolated from parasite cultures, digested into ribonucleosides and analyzed using liquid chromatography (LC) coupled to either a triple-quadrupole (QQQ) or a quadrupole time-of-flight (QTOF) mass spectrometer. Two approaches were combined, i.e. neutral loss scan (NLS) and multiple reaction monitoring (MRM) in LC-QQQ analysis. Identified ribonucleosides were validated by matching compound mass and retention time with synthetic standards and by collision-induced fragmentation (CID) analysis (except for mAm). b: Exemplary electropherograms of total RNA (top) and mRNA (bottom) from *P. falciparum.* The traces were generated by the Agilent 2100 Bioanalyzer and represent RNA concentrations (i.e. fluorescence intensity, y-axis) over mRNA length (nucleotides, [nt]). tRNA and 18S and 28S rRNA peaks are absent from the mRNA sample that was subjected to LC-MS/MS after two rounds of poly(A) RNA enrichment using oligo(dT) beads. c: Same as B for human mRNA extracted from A549 cells. d: LC-MS/MS extracted ion chromatograms (same as Fig. 1c), but only for m^6^A and at all time points sampled (top) and for all modified and canonical ribonucleosides (bottom, sampled at 36 hpi).

**Supplementary Fig. 2:**
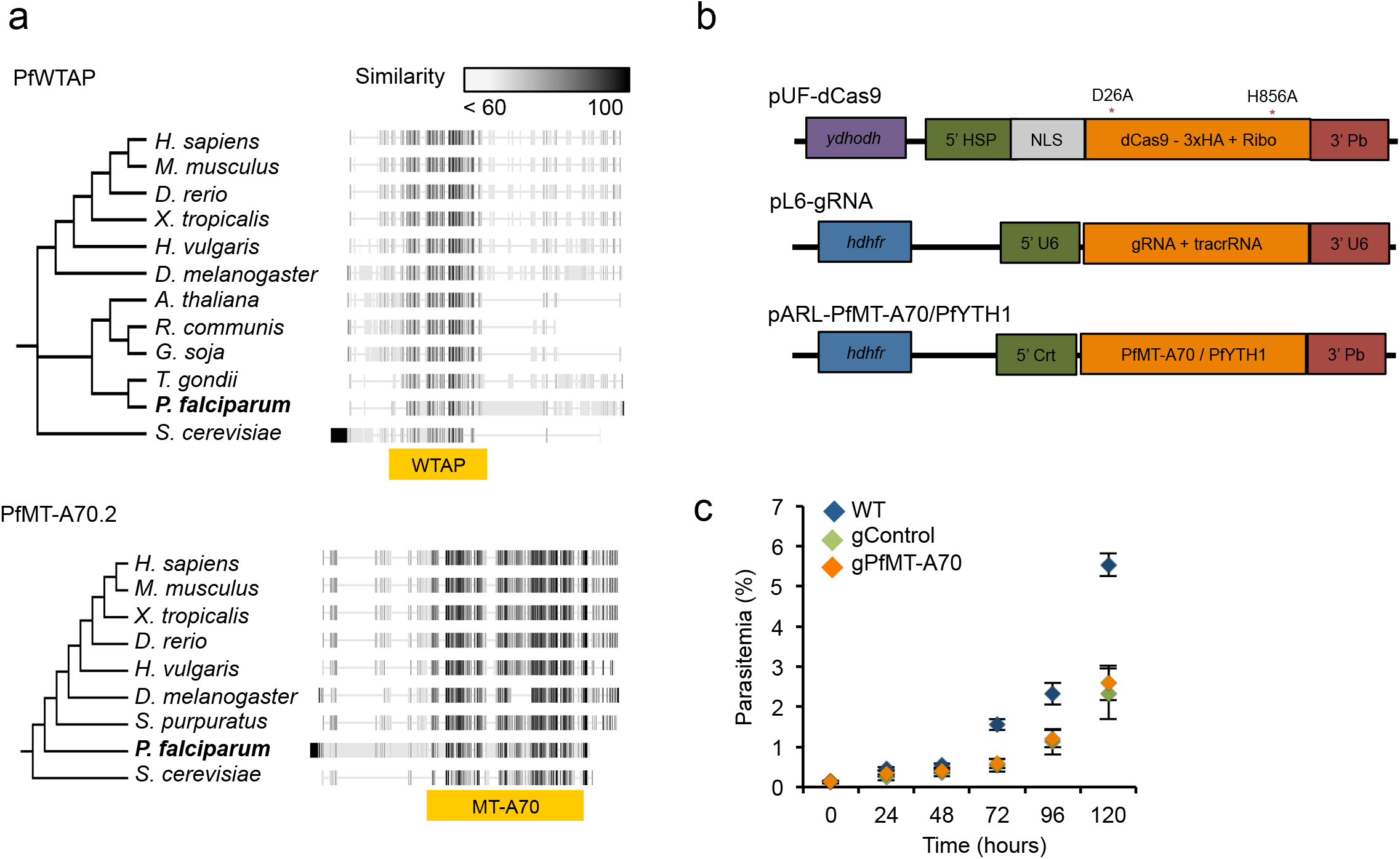
Identification of the m^6^A writer complex in *P. falciparum*. a: Phylogenetic trees and protein alignments of putative members of the *P. falciparum* core m^6^A writer complex that were co-immunoprecipitated with PfMT-A70: PfWATP (top) and PfMT-A70.2 (bottom). The locations of the characteristic domains are shown below the alignments (yellow bars). Grey scale indicates the percentage similarity across all proteins at an aligned position. b: Diagrams of plasmids used in this study. Top: pUF-dCas9 encoding the C-terminally HA-tagged dCas9, with catalytic domain mutations indicated with asterisks. Middle: pL6-gRNA encoding the guide and RNA that localize the dCas9 protein to the target site. Bottom: pARL plasmids for episomal expression of HA-tagged PfMT-A70 or PfYTH. Drug-selectable markers are indicated in purple for yeast dihydroorotate dehydrogenase (yDHODH) or blue for human dihydrofolate reductase (hDHFR). 5’ HSP: HSP90 promoter. NLS: nuclear localization signal. 5’ and 3’ U6: U6 snRNA polymerase III promoter and 3’ region, respectively. 5’ Crt: Chloroquine resistance transporter promoter. 3’ Pb: *P. berghei dhfr* 3’ region. c: Growth curve of wild-type (WT) parasites and parasites expressing dCas9 and the PfMT-A70 targeting (gPfMT-A70) or control (gControl) gRNA. Both dCas9 parasite cell lines show substantially reduced growth rates which can be attributed to the double drug selection pressure needed for plasmid maintenance. n = 6 (2 clones x 3 different blood samples). Error bars indicate standard error of the mean.

**Supplementary Fig. 3:**
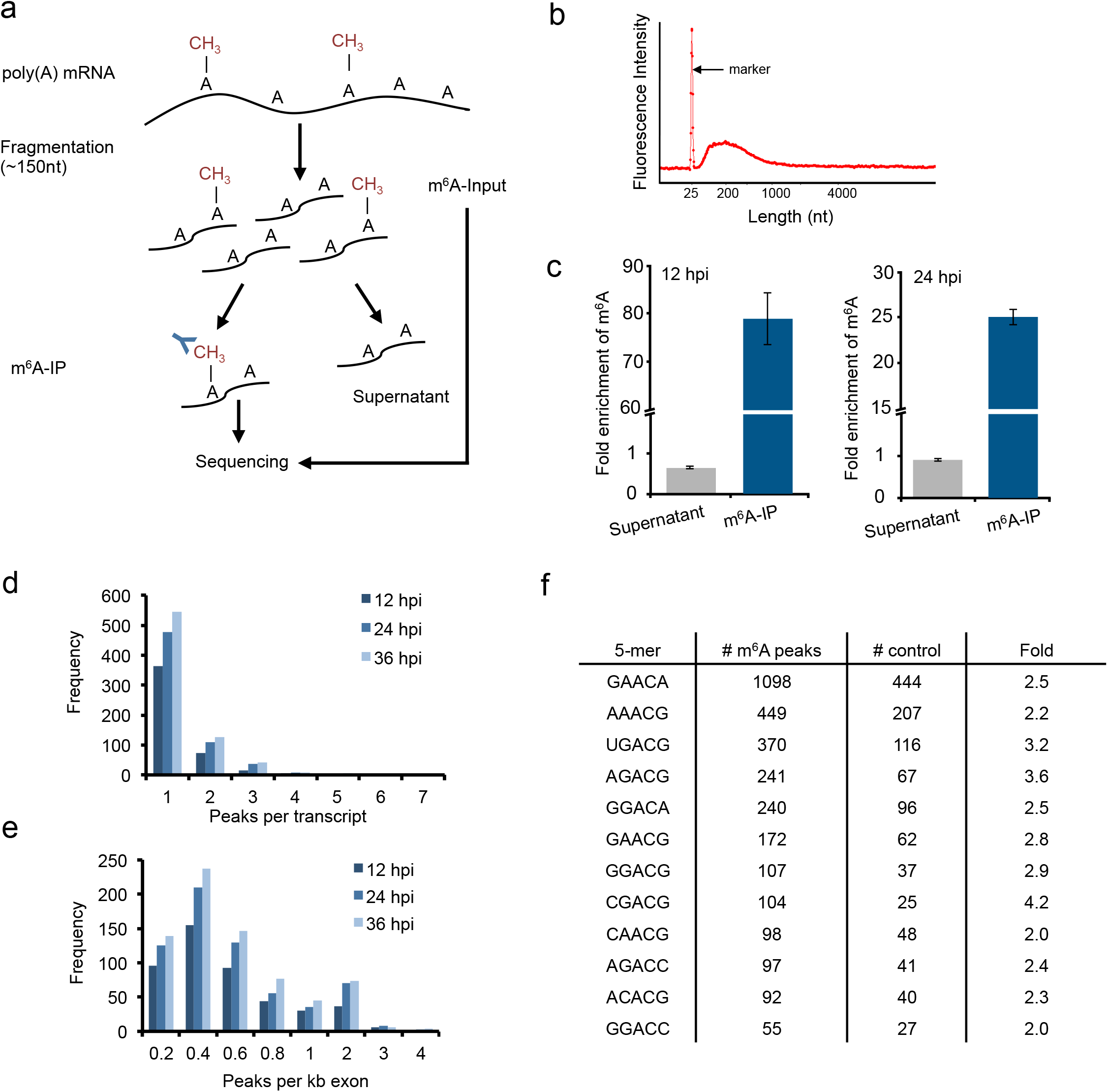
Identification of m^6^A methylation sites by m^6^A-seq. a: Simplified flowchart of the m^6^A-seq experiment. Poly-A enriched mRNA is fragmented to an average size of ~150 nt, and m^6^A-methylated fragments are immunoprecipitated using specific anti-m^6^A antibodies. Input controls and eluates are sequenced and used for m^6^A peak calling. b: Representative electropherogram of fragmented mRNA (see Supplementary Fig. Fig. 1b) used as input to the m^6^A-immunoprecipitation. The traces were generated by the Agilent 2100 Bioanalyzer and represent RNA concentrations (i.e. fluorescence intensity, y-axis) over mRNA length (nucleotides [nt], x-axis). The average fragment length is 150 nt. c: Fold enrichment of m^6^A levels as measured by LC-MS/MS in different fractions of the m^6^A-IP experiment at 12 and 24 hpi. m^6^A levels were normalized to the total number of adenosines (i.e. m^6^A/A) for each fraction and fold enrichments were calculated as ratio over the m^6^A/A levels in the input mRNA sample. Error bars indicate standard error of the mean for two technical replicates. d: Histogram showing the number of significant m^6^A peaks per transcript at the different time points of the IDC. e: Histogram showing the number of genes with different m^6^A peak densities (i.e. number of peaks per kilobase (kb) of exon) at the different time points of the IDC. f: Abundance of 5-mer sequences containing an XXACX (X = A/U/C/G) context in the m^6^A peak region (± 100 nt surrounding the peak summit) and random control sequences of similar size and length as well as the fold enrichment (Fold).

**Supplementary Fig. 4:**
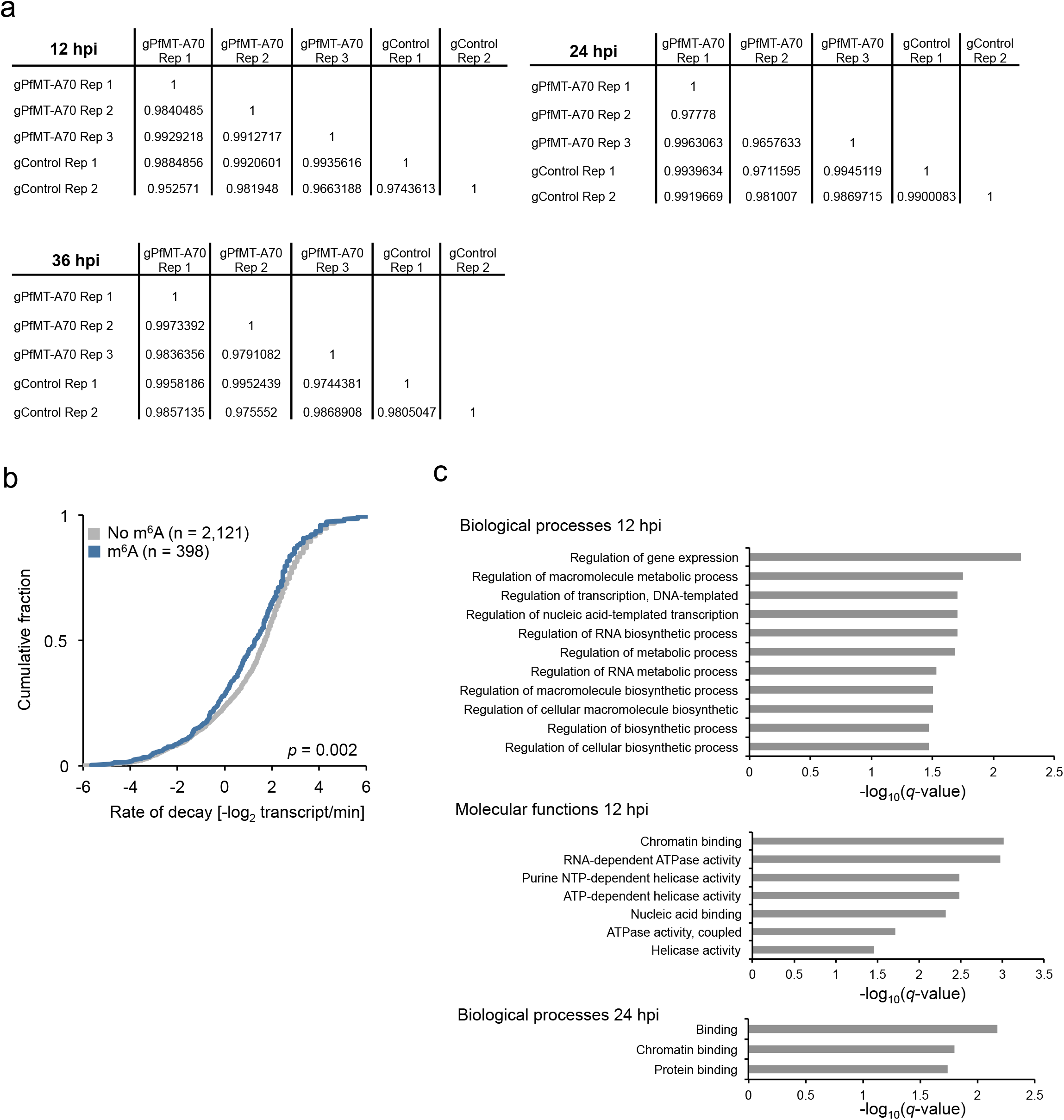
Analysis of differential expression and correlation of m^6^A with mRNA stability. a: Matrices summarizing the correlation (Pearson r) of gene counts in the three replicates of gPfMT-A70 and the two replicates of gControl at 12, 24 and 36 hpi. b: Comparison of mRNA stability (calculated by^13^) between m^6^A methylated (blue) and non-methylated (grey) transcripts. m^6^A methylated transcripts show significantly higher rates of mRNA degradation (transcripts/min) than non-methylated transcripts. c: Significantly enriched (FDR ≤ 0.05) Gene Ontology (GO) categories of ‘m^6^A-sensitive’ transcripts. Top: Enriched biological processes at 12 hpi. Middle: Enriched molecular functions at 12 hpi. Bottom: Enriched biological processes at 24 hpi.

**Supplementary Table 1**: Table of all detected mRNA modifications.

**Supplementary Table 2**: Calibration data for m^6^A/A quantification.

**Supplementary Table 3**: Summary table of the PfMT-A70 co-IP experiment

**Supplementary Table 4**: Summary Table containing annotation and enrichment of significant m^6^A sites detected at 12 hpi.

**Supplementary Table 5**: Summary Table containing annotation and enrichment of significant m^6^A sites detected at 24 hpi.

**Supplementary Table 6**: Summary Table containing annotation and enrichment of significant m^6^A sites detected at 36 hpi.

**Supplementary Table 7**: Summary of the differential expression analysis at 12, 24 and 36 hpi

**Supplementary Table 8**: Gene Ontology enrichment analysis of ‘m^6^A-sensitive’ transcripts at 12 and 24 hpi.

**Supplementary Table 9**: Table of primers used in this study

## Methods

### Parasite culture

Asexual blood stage *P. falciparum* parasites (strain 3D7) were cultured as described previously^4^. Briefly, parasites were cultured in human red blood cells (RBC) in RPMI-1640 culture medium (Thermo Fisher # 53400-025) supplemented with 10% v/v Albumax I (Thermo Fisher # 11020039), hypoxanthine (0.1 mM final concentration, CC-Pro # Z-41-M) and 10 mg gentamicin (Sigma # G1397-10ML) at 4% hematocrit and under 5% O_2_, 3% CO_2_ at 37 °C. Parasite development was monitored by Giemsa staining.

For synchronization, late stage parasites were enriched with plasmagel flotation followed by ring stage enrichment via sorbitol (5%) lysis 6 h later. For sampling of highly synchronous parasites during the IDC, the synchronous schizonts were enriched by plasmagel flotation shortly before reinvasion, followed by a sorbitol lysis 6 h later. The ‘zero’ hour time point was considered to be 3 h after plasmagel flotation. Parasites were collected at 4% hematocrit and ~2-3% parasitemia.

### Parasite growth assay

Parasite growth kinetics were measured as described previously^58^. Briefly, two clones of wild-type, gControl and gPfMT-A70 *P. falciparum* parasites were tightly synchronized and diluted to 0.2% parasitemia at ring stage (0 hours). The growth curve was replicated in three distinct batches of RBC and in triplicates on a 96 well plate (200 μl complete culture media per well). Parasitemia was measured every 24 hours by staining parasite nuclei using SYBR Green I (Sigma # S9430) and counting (un-) infected RBCs using a Guava easyCyte flow cytometer (Merck Millipore).

### Total RNA extraction

For RNA extraction, infected red blood cells (iRBCs) were collected by centrifugation and washed once with Dulbecco’s phosphate-buffered saline (DPBS) (Thermo Fisher # 14190-144) at 37 °C. iRBCs were lysed in 0.075 % saponin in DPBS at 37 °C, and the parasite cell pellet was washed once with DPBS and then resuspended in 700 μL QIAzol reagent (Qiagen # 79306). Total RNA was extracted using the miRNeasy Mini Kit (Qiagen # 217004), including an on-column DNase I digestion and exclusion of small RNA (< 200 nt) according to the manufacturer’s protocol.

### Analysis of mRNA modifications by LC-MS/MS

Total RNA from highly synchronous parasites was extracted from samples collected during the IDC at seven time points at six-hour intervals (i.e., 6 hpi to 42 hpi). To minimize human RNA and DNA contamination, parasites were cultured in leukocyte-depleted red blood cells. Total RNA from human A549 cells was extracted as described above. Poly(A) RNA from *P. falciparum* or human A549 total RNA was enriched by two successive rounds of purification with the Dynabeads mRNA purification kit (Thermo Fisher # 61006, Supplementary Fig. 1b,c).

Purified *P. falciparum* RNA was hydrolyzed enzymatically as described with a slightly modified protocol using the following components in the buffer mix [10 mM Tris-HCl (pH 7.9), 1 mM MgCb, 5 U Benzonase (Merck # 71206), 50 μM Desferroxamine (Sigma # D9533), 0.1 μg/μL Pentostatin (Sigma # SML0508), 100 μM Butylated hydroxytoluene (Sigma # W218405), 0.5 μg/μL Tetrahydrouridine (Calbiochem # 584222), 5 U Bacterial Alkaline Phosphatase (Thermo Fischer # 18011015), 0.05 U Phosphodiesterase I (Sigma # P3243)] and [^15^N]5-2’-deoxyadenosine ([^15^N]-dA) (Cambridge Isotope Laboratories # NLM-3895-25) as the internal standard to account for variations in sample handling and mass spectrometer response^59^. Hypersil GOLD aQ column [100 x 2.1 mm, 1.9 μm (Thermo Scientific # 25305)] was used to resolve the digested ribonucleosides in a two-buffer eluent system (buffer A = 0.1% formic acid in water and buffer B = 0.1% formic acid in acetonitrile). HPLC was performed at a flow rate of 300 μL/m at 25 °C. The gradient of 0.1% formic acid in acetonitrile was as follows: 0-12 min, held at 0%; 12-15.3 min, 0%-1%; 15.3-18.7 min, 1%-6%; 18.7-20 min, held at 6%; 20-24 min, 6%-100%; 24-27.3 min, held at 100%; 27.3-28 min, 100%-0%; 28-41 min, 0%. The HPLC column was directly connected to an Agilent 6490 triple quadrupole mass spectrometer or Agilent 6530 q-ToF with ESI Jetstream ionization operated in positive ion mode. The voltages and source gas parameters were as follows: gas temperature: 50 °C; gas flow: 11 L/min; nebulizer: 20 psi; sheath gas temperature: 300 °C; sheath gas flow: 12 L/min; capillary voltage 1800 V and nozzle voltage, 2000 V. The molecular transition ions were quantified in multiple-reaction monitoring (MRM) mode (Supplementary Table 1). For the collision-induced dissociation using the q-ToF, data was acquired from 200-500 m/z with an acquisition rate of 5 spectra/s in MS mode and from 50 to 500 m/z with an acquisition rate of 2 spectra/s in MS/MS mode.

### Generation of tagged PfMT-A70 and PfYTH *P. falciparum* strains

PfMT-A70 and PfYTH open reading frames (ORF) without stop codon were PCR-amplified from *P. falciparum* genomic DNA (strain 3D7) using primers MT-A70_F/MT-A70_R and YTH1_F/YTH1-R (Supplementary Table 9), respectively. A 3xFLAG-2xHA tag was synthesized by GenScript (Piscataway, USA) and PCR amplified using primer pair MT-A70_HA-F/HA_R for PfMT-A70 and YTH1_HA-F/HA_R for PfYTH (Supplementary Table 9). The resulting tag fragments were fused to the C-terminus of the corresponding ORF fragment with PCR using primer pairs MT-A70_F/HA_R for PfMT-A70 and YTH1_F/HA_R for PfYTH (Supplementary Table 9). The final PCR fragments were cloned into the XhoI/KpnI restriction sites of the pARL-STEVOR^full^ vector backbone^60^ using the In-Fusion HD cloning kit (Clontech # 639649). The resulting plasmids were verified by Sanger sequencing. All PCR reactions were performed using the KAPA HiFi DNA Polymerase (Roche # 07958846001) following the manufacturer’s protocol, but with lower elongation temperature (62°C or 68 °C). Cloning and plasmid amplification were performed using XL10-Gold ultracompetent *E. coli* (Agilent Technologies # 200315) following the manufacturer’s protocol. The final plasmids (pARL-PfMT-A70-3xFLAG-2xHA and pARL-PfYTH-3xFLAG-2xHA) contain the drug-selectable marker human dihydrofolate reductase (hDHFR), conferring resistance to the antifolate drug WR99210. The final plasmids encode a full length PfMT-A70 or PfYTH protein with a C-terminal 3xFLAG-2xHA tag under the chloroquine resistance transporter promoter (Fig. S2B).

Ring stage parasites were transfected with 50 μg plasmid by electroporation as described previously^61^ and drug-resistant parasites emerged after four weeks of continuous culture in the presence of 2.67 nM WR99210 (Jacobus Pharmaceuticals).

### Protein fractionation and Western Blot analysis

One mL iRBCs containing synchronous parasites expressing HA-tagged PfMT-A70 or PfYTH at ring stage (2-3% parasitemia) were washed once with DPBS at 37°C, and RBCs were lysed with 0.075% saponin in DPBS. Parasites were then washed once again with DPBS. For separation of the cytoplasmic and nuclear protein fractions, the cell pellet was first resuspended in one mL cytoplasmic lysis buffer (CLB: 25 mM Tris-HCl [pH 7.5], 10 mM NaCl, 1% IGEPAL CA-630, 1mM DTT, 1.5mM MgCl_2_, 1X Protease inhibitor [PI, Roche # 11836170001]) and incubated on ice for 30 min. Cells were further homogenized in a glass douncer, and the cytoplasmic lysate was cleared with centrifugation (13,500 *g*, 10 min, 4 °C). The pellet (containing the nuclei) was resuspended in 100 μL nuclear extraction buffer (25 mM Tris-HCl [pH 7.5], 1mM DTT, 1.5mM MgCl_2_, 600 mM NaCl, 1% IGEPAL CA-630, PI) supplemented with 1 μL Benzonase (Sigma Aldrich # E1014-5KU) and sonicated for 10 cycles with 30 s (on/off) intervals (5 min total sonication time) in a Diagenode Bioruptor Pico (Diagenode # B01060010). This nuclear lysate was cleared with centrifugation (13,500 g, 10 min, 4°C), and the nuclear fraction was diluted with 300 μL CLB.

Protein samples were supplemented with NuPage Sample Buffer (Thermo Fisher # NP0008) and NuPage Reducing Agent (Thermo Fisher # NP0004) and denatured for 10 min at 70 °C. The samples were separated on a NuPage 4-12% Bis-Tris gel using MOPS running buffer at 150 V for 1.5 h and transferred to a PVDF membrane overnight at 15 V at 4 °C. The membrane was blocked for 1 h in 1% milk in 0.1% Tween20 in PBS (PBST). The HA-tagged proteins and Histone H3 were detected with anti-HA (Abcam # ab9110; 1:1000 in 1% milk-PBST) and anti-H3 (Abcam # ab1791: 1:1000 in 1% milk-PBST) primary antibodies, respectively, followed by donkey anti-rabbit (GE # NA934-1ML) secondary antibodies conjugated to HRP (1:5000). PfAldolase was detected using an HRP conjugated anti-PfAldolase (Abcam # ab38905, 1:5000 in 1% milk PBST) antibody. The HRP signal was developed using the SuperSignal West Pico chemiluminescent substrate (Thermo Fisher # 34580) and imaged with a ChemiDoc XRS+ (Bio-Rad).

### PfMT-A70 co-immunoprecipitation

*P. falciparum* PfMT-A70-HA (*n* = 3 replicates) expressing parasites (cultured with 2.67 nM WR99210 [Jacobus Pharmaceuticals]) and wild-type *P. falciparum* 3D7 (i.e. as negative control, cultured in standard growth medium) were sorbitol-synchronized After 36 hours, the parasites were harvested after Percoll (Sigma # P4937) enrichment, washed twice with RPMI and crosslinked with 0.5 mM dithiobissuccinimidyl propionate (DSP) (Thermo Fisher # 22585) in PBS for 60 min at 37°C^62^. The reaction was quenched with PBS containing 25 mM Tris-HCl. These trophozoites were lysed with RIPA buffer (10 mM Tris HCl pH 7.5, 150 mM NaCl, 0.1% SDS, 1% Triton) containing protease and phosphatase inhibitor cocktail (Thermo Fisher # 78440). The lysates were cleared by centrifugation at 16,000 g for 10 min. Supernatants were incubated with 25 μl of anti-HA Dynabeads (Thermo Fisher # 88836) overnight at 4°C. Beads were magnetically isolated and washed five times with 1mL RIPA buffer following five washes with 1mL PBS and one wash with 1 mL 100 mM Ammonium Bicarbonate (Sigma # 09830). The beads were reduced with 10 mM dithiothreitol (Sigma # D9779), alkylated with 55 mM iodoacetamide (Sigma # I1149) and subjected to on-bead digestion using 50 μg of trypsin (Thermo Fisher # 90059). The resulting peptides were desalted using C18 cartridges (Thermo Fisher # 89852) and sent for MS-analysis.

### Protein mass-spectrometry

Peptides were separated by reverse phase HPLC (Thermo Easy nLC1000) using a precolumn (made in house, 6 cm of 10 μm C18) and a self-pack 5 μm tip analytical column (12 cm of 5 μm C18, New Objective) over a 150 minute gradient before nanoelectrospray using a Q-Exactive HF-X mass spectrometer (Thermo Scientific). The mass spectrometer was operated in a data-dependent mode. The parameters for the full scan MS were: resolution of 70,000 across 350-2000 *m/z,* AGC 3e^6^, and maximum IT 50 ms. The full MS scan was followed by MS/MS for the top 15 precursor ions in each cycle with a NCE of 28 and dynamic exclusion of 30 s. Raw mass spectral data files (.raw) were searched using Proteome Discoverer (Thermo Fisher) and Mascot (version 2.4.1)^63^. Mascot search parameters were: 10 ppm mass tolerance for precursor ions; 15 mmu for fragment ion mass tolerance; 2 missed cleavages of trypsin; fixed modification was carbamidomethylation of cysteine; variable modifications were methionine oxidation and serine, threonine and tyrosine phosphorylation. Only peptides with a Mascot score greater than or equal to 25 and an isolation interference less than or equal to 30 were included in the data analysis.

### CRISPR interference knockdown of PfMT-A70

An enzymatically inactive or ‘dead’ Cas9 protein (dCas9) was expressed from the pUF-dCas9 plasmid, which is derived from the pUF-Cas9 plasmid backbone described in^61^ with the following modifications: the Cas9 was mutated at the RuvC and HNH positions, as described in ref. 43, and the 3xFLAG tag present at the N-terminus of the Cas9 protein in pUF-Cas9 was removed and a 3xHA tag followed by a *glmS* ribozyme was added to the C-terminus of the dCas9 protein in pUF-dCas9 (Supplementary Fig. 2b). Candidate target sites for dCas9 had to be 1) specific, 2) close to the transcription start site, and 3) on the non-template strand and were identified using ‘Protospacer’^64^.

The guide RNA targeting the PfMT-A70 promoter (gPfMT-A70) (Supplementary Table 9) was synthesized by Eurofins Genomics (Ebersberg, Germany) and cloned into the BtgZ1 restriction site of the pL6-egfp plasmid as described elsewhere^61^ using the In-Fusion HD cloning kit (Clontech # 639649). For the gControl gRNA, the pL6-egfp plasmid was unaltered. Cloning and plasmid amplification were performed using XL10-Gold ultracompetent *E. coli* (Agilent Technologies # 200315) following the manufacturer’s protocol and plasmids were purified using the NuceloBond Xtra Maxi Plus kit (Macherey Nagel # 740416.10) according to the manufacturer’s instructions. Ring stage parasites were transfected with 25 μg of pL6 and pUF plasmids by electroporation as described previously^61^ and drug-resistant parasites emerged after four weeks of continuous culture in the presence of 2.67 nM WR99210 (Jacobus Pharmaceuticals) and 1.5 μM DSM1. Parasite cloning was performed by limiting dilution as described previously^61^.

### Quantitative reverse-transcription PCR

Total RNA was extracted from gPfMT-A70 and gControl parasites collected at 12, 24, and 36 hpi, as described above. cDNA was generated using the Superscipt VILO cDNA synthesis kit (Thermo Fisher # 11754050) following the manufacturer’s protocol. PfMT-A70 was amplified in technical triplicates using Power Sybr Green PCR Master Mix (Thermo Fisher # 4367659) and primers PfMT-A70_qPCR-F/PfMT-A70_qPCR-R (Supplementary Table 9) on a BioRad CFX qPCR machine. A no RT control (substitution of RT during cDNA synthesis with H_2_O) and a no template control (no cDNA added during qPCR amplification) were included in all experiments. Mean starting quantity (SQ-mean) values of technical triplicates were extrapolated from a standard curve made with serial dilutions of genomic DNA. PfMT-A70 levels were normalized to those of an internal housekeeping control gene serine-tRNA ligase (PF3D7_0717700) using primer pair Seryl_qPCR-F/Seryl_qPCR-R (Tabe S7).

### dCas9 chromatin immunoprecipitation (dCas9 ChIP)

ChIP experiments were performed for gPfMT-A70 and gControl parasites at 12 hpi at ~2% parasitemia. Red blood cell cultures (~4% hematocrit) were crosslinked for 10 min by direct addition of methanol-free formaldehyde (1% final concentration) (Thermo Fisher # 28908). The cross-linking reaction was quenched using a final concentration of 0.125 M Glycine for an additional 5 min at room temperature. The iRBC pellet was washed once with DPBS at 4C and lysed using 0.15% saponin in DPBS at 4 °C. Parasite cells were collected by centrifugation (3250 g, 4 °C, 5 min), washed twice in DPBS at 4°C, snap frozen in liquid nitrogen, and stored until further use at −80 °C.

For each ChIP experiment, ~2 x 10^8^ parasites were resuspended in 2 mL cytoplasmic lysis buffer (10 mM HEPES [pH 8], 10 mM KCl, 0.1 mM EDTA [pH 8], plus complete protease inhibitor [PI, Roche # 11836170001]), transferred to a prechilled 2 mL douncer homogenizer and set on ice for 30 min. 50 μL 10% IGEPAL CA-630 were added (0.25% final concentration) and parasites were subjected to dounce homogenization. The nuclei were pelleted by centrifugation (10 min, 13500 g, 4 °C), and the supernatant (cytoplasmic fraction) was removed. The nuclei pellet was resuspend in 300 μL SDS Lysis buffer (50 mM Tris-HCl [pH 8], 10 mM EDTA [pH 8], 1% SDS, PI) and transferred to a Diagenode 1.5 mL sonication tube. Chromatin was sonicated to ~ 1,000 bp DNA fragments in a Bioruptor Pico Plus for 24 cycles with 30 s (on/off) intervals at high intensity (12 min total sonication time), and remaining debris were removed by centrifugation (10 min at 13,500 *g* at 4 °C). 30 μL of sample were stored as input control at −80 °C. 120 μL of sample were diluted 1:10 in ChIP dilution buffer (16.7 mM Tris-HCl [pH 8], 150 mM NaCl, 1.2 mM EDTA [pH 8], 1% Triton X-100, 0.01% SDS, PI).

For each sample, 25 μL of Protein G magnetic beads (Thermo Fisher # 10004D) were used and pre-washed three times in 500 μL ChIP dilution buffer. Beads were resuspended in 100 μL ChIP dilution buffer and incubated with 1 μg anti-HA (Abcam # ab9110) antibody for 2 h with constant rotation at 4 °C. Following antibody binding, beads were washed twice in 1 mL ChIP dilution buffer and resuspended in 25 μL ChIP dilution buffer. The ChIP sample was added to the antibody-conjugated beads and incubated overnight at 4°C with constant rotation. Following immunoprecipitation, the beads were washed sequentially in 1 mL of low salt wash buffer (20 mM Tris-HCl [pH 8], 150 mM NaCl, 2 mM EDTA [pH 8], 1% Triton X-100, 0.1% SDS, PI), 1 mL high salt wash buffer (20 mM Tris-HCl [pH 8], 500 mM NaCl, 2 mM EDTA [pH 8], 1% Triton X-100, 0.1% SDS, PI), one mL LiCl wash buffer (10 mM Tris-HCl [pH 8], 250 mM NaCl, 1 mM EDTA [pH 8], 0.5% IGEPAL CA-630, 0.5% sodium deoxycholate, PI) and 1 mL TE buffer (10 mM Tris-HCl [pH 8], 1 mM EDTA [pH 8]). For each wash, the beads were rotated for 5 min at 4°C except for the last TE wash, which was performed at room temperature. The beads were subsequently resuspended in 210 μL elution buffer (50 mM Tris-HCl [pH 8], 10 mM EDTA [pH 8], 1% SDS) and protein-DNA complexes were eluted by 30 min incubation at 65°C in an agitator (900 rpm 1 min, still 30 s). At this step, the input sample was recovered from storage, diluted with 170 μL elution buffer, and processed in parallel with the IP sample.

Protein-DNA complexes were reverse crosslinked by incubation at 65°C for 6h. Samples were diluted with 200 μL of TE buffer and incubated with RNaseA (0.2 mg/ml final concentration) (Thermo Fisher # EN0531) at 37°C for 2 h. Proteinase K (0.2 μg/ml final concentration) (NEB # P8107S) was then added and samples were incubated at 55°C for two hr. DNA was purified by adding 400 μl phenol:chloroform:isoamyl alcohol, and phases were separated by centrifugation (10 min, 13,500 g, 4 °C) after vigorous vortexing. 16 μL of 5 M NaCl (200 mM final concentration) and 30 μg glycogen were added to the aqueous phase, and DNA was precipitated by adding 800 μl 100% EtOH 4°C and incubating for 30 min at −20 °C. DNA was pelleted by centrifugation (20,000 g, 10 min, 4 °C) and washed with 500 μL 80% EtOH 4 °C. After centrifugation, the DNA pellet was air-dried and resuspended in 30 μL of 10 mM Tris-HCl, pH 8.0.

### m^6^A immunoprecipitation and sequencing

Two replicates were sampled for both cell lines expressing dCas9 and either a non-targeting control gRNA (gControl) or the PfMT-A70 promoter-targeting gRNA (gPfMT-A70). Total RNA from highly synchronous parasites was collected at 12, 24, and 36 hpi as described above, followed by poly(A) RNA enrichment using the Dynabeads mRNA purification kit (Thermo Fisher # 61006).

300-500 ng purified mRNA was fragmented for 15 min at 70°C to ~100 nt using the NEBNext fragmentation module (NEB # E6150S) and purified using the Qiagen MinElute kit (Qiagen # 74204) with slight modifications: To also purify small RNA < 200 nt, 100 μL of RNA sample was supplemented with 350 μL of lysis buffer (provided in the kit) followed by the addition of 700 μL of 100% ethanol. 20 ng fragmented mRNA were stored as input sample at −80 °C.

For each m^6^A-IP, 25 μL of Protein A/G magnetic beads (Thermo Fischer # 88802) were diluted in 250 μL reaction buffer (150 mM NaCl, 10 mM Tris-HCl, pH 7.5, 0.1% IGEPAL CA-630 in nuclease-free H_2_O) and washed twice in 250 μL reaction buffer. The beads were resuspended in 100 μL reaction buffer and incubated with 2.5 μg anti-m^6^A antibody (Abcam # ab151230) for 30 min at room temperature with constant rotation. The beads were subsequently washed three times with 0.5 mL reaction buffer. For each m^6^A-IP, 300-500ng fragmented mRNA were diluted with nuclease-free H_2_O to a final volume of 395 μL, supplemented with 5 μL RNase inhibitor (Promega # N2615), 100 μL 5X reaction buffer (750 mM NaCl, 50 mM Tris-HCl, pH 7.5, 0.5% IGEPAL CA-630 in nuclease-free H_2_O) and added to the antibody-bead mixture. The sample was incubated at 4°C with constant rotation for 2 h. Following immunoprecipitation, the supernatant was removed and the beads were washed twice in 250 μL reaction buffer, twice in 250 uL low salt wash buffer (50 mM NaCl, 10 mM Tris-HCl, pH 7.5, 0.1% IGEPAL CA-630 in nuclease-free H_2_O), and twice in 250 μL high salt wash buffer (500 mM NaCl, 10 mM Tris-HCl, pH 7.5, 0.1% IGEPAL CA-630 in nuclease-free H_2_O), all at 4 °C. m^6^A mRNA fragments were competitively eluted by two successive incubations (1 h, 1000 rpm continuous shaking at 4 °C) with 100 μL elution buffer (6.7 mM m^6^A 5-monophosphate sodium salt [Sigma # M2780-10MG], 150 mM NaCl, 10 mM Tris-HCl, pH 7.5, 0.1%, 400 U RNase inhibitor [Promega # N2615] in nuclease-free H_2_O). RNA was purified using the Qiagen MinElute kit (Qiagen # 74204) as described above.

RNA sequencing libraries were prepared using the Illumina TruSeq stranded RNA Library Prep Kit (Illumina # RS-122-2101) according to the manufacturer’s instructions with slight modifications. To account for the AT-richness of cDNA fragments, we used the KAPA Hifi polymerase (Roche # 07958846001) at the library amplification step. The libraries were sequenced on an Illumina NextSeq 500 platform with a 1×150 bp single-end read layout.

### RNA-sequencing for differential gene expression analysis

For RNA sequencing, three replicates of highly synchronous gPfMT-A70 and two replicates of gControl parasites were collected at 12, 24 and 36 hpi. Total RNA was extracted as described above and poly(A) RNA was enriched using the Dynabeads mRNA purification kit (Thermo Fisher # 61006).

RNA sequencing libraries were prepared using the Illumina TruSeq stranded RNA Library Prep Kit (Illumina # RS-122-2101) according to the manufacturer’s instructions with slight modifications. To account for the AT-richness of cDNA fragments, we used the KAPA Hifi polymerase (Roche # 07958846001) at the library amplification step. The libraries were sequenced on an Illumina NextSeq 500 platform with a 1×150 bp single-end read layout.

### Phylogenetic analysis of the m^6^A writer and reader complex

Candidate proteins of the *P. falciparum* m^6^A writer complex were identified by searching known members of the m^6^A writer complex from other eukaryotes against the *P. falciparum* proteome using blastp^65^. Protein alignments were generated using mafft^66^ and visualized in Geneious^67^. For the phylogenetic analysis, gaps in the protein alignments were removed with trimal^68^ and the best phylogenetic model was calculated using protTest3^69^. Phylogenetic trees were constructed using MEGA (v7)^70^ with 1,000 bootstrap replicates, and the bootstrap consensus trees were visualized in FigTree (v1.4.3, http://tree.bio.ed.ac.uk/software/figtree/).

### Identification and quantification of mRNA modifications by LC-MS/MS

LC/MS data was extracted using the MassHunter Qualitative and Quantitative Analysis Software (version B06.00) and further analyzed as follows. Briefly, to account for any background signal that could be a contribution from the salts and enzymes in the digestion buffer, the signal intensity (i.e. area under the curve) for each ribonucleoside is first subtracted from a matrix sample (without any RNA). To calculate relative levels of each modified ribonucleoside and to adjust for different injection amounts of RNA in each sample, the matrix-corrected intensity of the modified ribonucleosides (e.g. m^6^A) was then divided by the intensity of the respective canonical ribonucleoside (e.g. rA). To adjust day-to-day fluctuation in retention time of the mass spectrometer, an internal standard ([15N]-2’-deoxyadenosine) was spiked into each sample during RNA digestion.

The absolute values for N^6^-methyladenosine (Fig. 1E) were calculated by using an external calibration curve prepared using known standards to account for the difference in ionization efficiency in the mass spectrometer (Supplementary Table 2).

For visualization of lifecycle-dependent changes in the quantities of modified ribonucleosides (Fig. 1b), the modified ribonucleoside ratios (i.e. m^6^A/A) were scaled to row Z-sores following the formula z = (x-u)/s, with x = actual measured value; u = average over all samples and s = standard deviation over all samples.

### dCas9 ChIP sequencing and analysis

DNA sequencing libraries were prepared using the Diagenode MicroPlex Library Preparation kit (Diagenode # C05010014) according to the manufacturer’s instructions and sequenced on an Illumina NextSeq 500 platform with a 1×150 bp single-end read layout. Raw image files were converted to fastq sequence files using Illumina’s bcl2fastq (v2.19). Reads were aligned to the *P. falciparum* genome (plasmoDB.org, v3, release v28)^35,71^ using bwa ‘mem’^72^ allowing only unique read mappings (option ‘-c 1’). Optical duplicates and alignments with a quality score < 20 were removed using samtools’ ‘rmdup’ and ‘view’^73^, respectively.

For coverage plots of gPfMT-A70 and gControl (Fig. 3b) ChIP-seq experiments, deeptool’s bamCompare^74^ was used to normalize the read coverage per genome position (option ‘--bs 1’) in the respective input and ChIP samples to the total number of mapped reads in each library (option ‘--normalizeUsingRPKM’). Normalized input coverage per bin was subtracted from the ChIP values (option ‘--ratio subtract’) and coverage plots were visualized in plasmoDB’s build-in genome browser together with other chromatin features (plasmoDB.org).

For genome-wide coverage plots (Fig. 3c), the same approach was used as above, but genome coverage was calculated not per genome position but for 1000 nt windows. To further remove background levels of unspecific dCas9 binding, the gControl ChIP-input coverage was subtracted from the gPfMT-A70 ChIP-input coverage and visualized using circos^75^.

### RNA sequencing read mapping

Raw image files were converted to fastq sequence files using Illumina’s bcl2fastq (v2.19). The reads were aligned to the *P. falciparum* genome (plasmoDB.org, v3, release 28)^35^ using bwa ‘mem’^72^ allowing only unique read mappings (option ‘-c 1’). Optical duplicates and alignments with a quality score < 20 were removed using samtools’ ‘rmdup’ and ‘view’^73^, respectively.

### m^6^A peak calling

At each time point, we separately identified significant (FDR ≤ 0.05) m^6^A peaks for each replicate (i.e. 2x gControl and 2x gPfMT-A70) using macs2^76^ with default settings but without prior peak modeling (option ‘--nomodel’) and the fragment size set to 150 bp (option ‘--extsize 150’). To generate a single robust set of m^6^A peaks for each time point, we identified overlapping enriched regions using bedtools ‘multiinter’^77^ and only retained those that were present in ≥ two of the four samples at each time point. Overlaps of m^6^A peaks and genic regions were identified using bedtools ‘intersect’^77^.

### m^6^A motif identification

For the m^6^A motif analysis, sequences ± 100 bp of the m^6^A peak summit were extracted from the *P. falciparum* genome using bedtools ‘getfasta’^77^, taking the strandedness of the underlying transcript into account (option ‘-s’). A random set of control sequences was constructed as following. For each m^6^A site in a transcript, a random position in the same transcript was calculated using bash ‘shuffle’ implemented in a custom script. Sequences ± 100 bp surrounding the random site were extracted using bedtools ‘getfasta’ as described above. Significantly enriched motifs in the m^6^A peak region were subsequently identified with homer2 ‘findMotif.pl’^78^.

5-mers in the m^6^A peak region and controls sequence set were counted using jellyfish^79^. The deduced sequence motif was generated using all 5-mers with an AC context that were enriched ≥ two-fold in the m^6^A peak region compared to the random control using weblogo^80^, taking their frequency in the m^6^A peak region into account. The central enrichment of the deduced sequence motif was calculated using homer2 ‘annotatePeaks.pl’.

### m^6^A enrichment calculation

m^6^A enrichments were calculated following the approach reported in ref. 51 and ref. 81. Briefly, for each time point, the number of reads mapping to an m^6^A peak in the m^6^A-IP and m^6^A-input sample was calculated using bedtools ‘coverage-count’ and normalized to the total number of mapped reads in each sample. m^6^A enrichments were then calculated as m^6^A-IP/input for each peak and in each individual sample. To calculate decreases in global m^6^A enrichment in gPfMT-A70 (Fig. 5b) at each time point, m^6^A enrichments of every peak were averaged over the two replicates of each condition. Of note, m^6^A enrichments were highly reproducible (Fig. 5b) and the individual m^6^A enrichment values of each m^6^A peak and replicate are shown in Supplementary Table 4-6).

For visualization of changes in m^6^A enrichments (Fig. 5a), we used cgat’s ‘bam2geneprofile’^82^. Briefly, the read coverage from m^6^A-IP and m^6^A-input libraries of each sample was calculated and normalized to the total size of mapped reads in each library in 10 nt windows and in a region ± 2 kb surrounding the m^6^A peak summit. m^6^A enrichments in each 10 nt window were calculated as the ration of m^6^A-IP over m^6^A-input.

### Differential gene expression analysis

RNA sequencing reads for three replicates of the gPfMT-A70 and two replicates of the gControl cell lines were mapped to the *P. falciparum* genome and quality filtered as described above. Strand specific gene counts were calculated using htseq-count^83^.

Differential gene expression analysis was performed using DESeq2^84^ using all replicates of gPfMT-A70 and gControl at each time point. Due to their monoallelic nature, genes encoding variant surface antigens (i.e. *var, rifin, stevor)* were excluded from the analysis. MA-plots were generated using the ‘baseMean’ (i.e. mean normalized read count over all replicates and conditions) and ‘log_2_FoldChange’ values (i.e. gPfMT-A70 over gControl) as determined by DESeq2 for each time point.

For the comparison of transcript abundances (Fig. 5c, left), we first averaged the m^6^A enrichment of every m^6^A peak over the two replicates of each condition as described above. We then retained only those m^6^A-methylated transcripts that harbor m^6^A peaks with the most pronounced decreases in m^6^A enrichment (i.e. ≥ two-fold decrease). For the direct comparison of transcript log_2_ fold changes (Fig. 5c, right), the set of transcripts with reduced m^6^A enrichments was compared to non-methylated transcripts being expressed in the same range of mean normalized read counts (i.e. baseMean value).

RPKM (reads per kilobase per exon per one million mapped reads) values were calculated in R using the command rpkm() from the package edgeR^85^.

### mRNA stability and translational efficiency analysis

For the comparative analysis of m^6^A methylation and mRNA stability or translation efficiency, we used m^6^A-methylated transcripts that were expressed in the top two quartiles of all transcripts at a given time point as calculated by RNA-seq of the m^6^A-input samples and with a peak density of ≥ 0.2 m^6^A sites per kilobase of exon. Corresponding mRNA stability (measured as mRNA half-life) and translational efficiency values of the same parasite developmental stages were extracted from^14^ and^17^, respectively. Independent measurements of mRNA stability data were retrieved from^13^.

### Statistical analysis

All statistical analysis were performed in R^86^. To test for a normal distribution of the data, the Shapiro-Wilk normality test was used. To test for significance between two groups, a two-tailed independent-samples *t*-test or Mann-Whitney U test were performed as indicated. Gene Ontology enrichments were calculated using the build-in tool at plasmoDB.org. Correlations between replicates were calculated in R using function cor() with default settings (i.e. calculation of Pearson correlation coefficient r).

All heatmaps were visualized using the function heatmap2() in the R gplots package with row z-score normalization. Z-scores were calculated over all samples (i.e. ‘rows’) following the formula z = (x-u)/s, with x = actual measured value; u = average over all samples and s = standard deviation over all samples.

### Data and availability

All data are accessible on the Gene Expression Omnibus database (https://www.ncbi.nlm.nih.gov/geo) under study accession number GSE123839. Raw sequence data are accessible on the NCBI Sequence Read Archive (SRA) under BioProject accession number PRJNA473770.

